# Neutrality in the Metaorganism

**DOI:** 10.1101/367243

**Authors:** Michael Sieber, Lucía Pita, Nancy Weiland-Bräuer, Philipp Dirksen, Jun Wang, Benedikt Mortzfeld, Sören Franzenburg, Ruth A. Schmitz, John F. Baines, Sebastian Fraune, Ute Hentschel, Hinrich Schulenburg, Thomas C. G. Bosch, Arne Traulsen

**Affiliations:** Max Planck Institute for Evolutionary Biology, Plön, Germany; GEOMAR Helmholtz Centre for Ocean Research Kiel, Kiel, Germany; Institute for General Microbiology, Christian-Albrechts-University Kiel, Kiel, Germany; Zoological Institute, Christian-Albrechts-University Kiel, Kiel, Germany; Institute of Microbiology, Chinese Academy of Science, Beijing, China; Institute for Clinical Molecular Biology, Christian-Albrechts-University Kiel, Kiel, Germany; Institute for Experimental Medicine, Christian-Albrechts-University Kiel, Kiel, Germany

## Abstract

Almost all animals and plants are inhabited by diverse communities of microorganisms, the microbiota, thereby forming an integrated entity, the metaorganism. Natural selection should favor hosts that shape the community composition of these microbes to promote a beneficial host-microbe symbiosis. Indeed, animal hosts often pose selective environments, which only a subset of the environmentally available microbes are able to colonize. How these microbes assemble after colonization to form the complex microbiota is less clear. Neutral models are based on the assumption that the alternatives in microbiota community composition are selectively equivalent and thus entirely shaped by random population dynamics and dispersal. Here, we use the neutral model as a null hypothesis to assess microbiata composition in host organisms, which does not rely on invoking any adaptive processes underlying microbial community assembly. We show that the overall microbiota community structure from a wide range of host organisms, in particular including previously understudied invertebrates, is in many cases consistent with neutral expectations. Our approach allows to identify individual microbes that are deviating from the neutral expectation and which are therefore interesting candidates for further study. Moreover, using simulated communities we demonstrate that transient community states may play a role in the deviations from the neutral expectation. Our findings highlight that the consideration of neutral processes and temporal changes in community composition are critical for an in-depth understanding of microbiota-host interactions.

---

The microbial communities living in and on animals can affect many important host functions, including metabolism [1, 2, 3], the immune system [4, 5, 6], and even behaviour [7, 8]. The extent and direction of this microbe-mediated influence is often linked to the presence or absence of species and their relative abundances. It is thus paramount to understand how host-associated microbial communities are assembled. A classical explanation for the emergence of a particular ecological community structure posits that every species is defined by distinct traits and occupies a specific ecological niche. An implicit hypothesis underlying much of microbiome research is that hosts have the potential to actively shape their associated microbial communities by providing niches for useful microbes [9]. This implies that individual hosts could select for a potentially very specific community structure.

While the assumption that the metaorganism – the host together with its associated microbes [10, 11] – is an actively shaped symbiotic unit is appealing, it may bias interpretation of microbiota-host analyses. An example is the widely reported connection between the structure of the human gut microbiota and obesity, which turns out to be very hard to distinguish from random noise [12]. More recently it has also been demonstrated that the genetic background of the host does not significantly shape human microbiome composition [13]. Indeed, it has been argued that selective processes should not form the null hypothesis for explaining the allegedly cooperative host-microbe symbiosis [14]. Instead, neutral models have been proposed as a valuable tool for finding patterns in the tremendous complexity of ecological communities that may have a deeper mechanistic cause. Neutral models assume ecological equivalence between species. Thus, the community structure within a single host is the outcome of purely stochastic population dynamics, immigration and local extinctions [15]. The Unified Neutral Theory of Biodiversity [16, 17] extends the neutral framework from a single site to multiple sites, each harbouring its own local community (Fig 1). Diversity within the local communities is maintained by immigration from a common source community, so that the whole setup resembles a mainland-island structure [18]. Neutral theory has been applied to numerous ecological systems and sparked a lot of controversy along the way [19].

**Figure 1.**
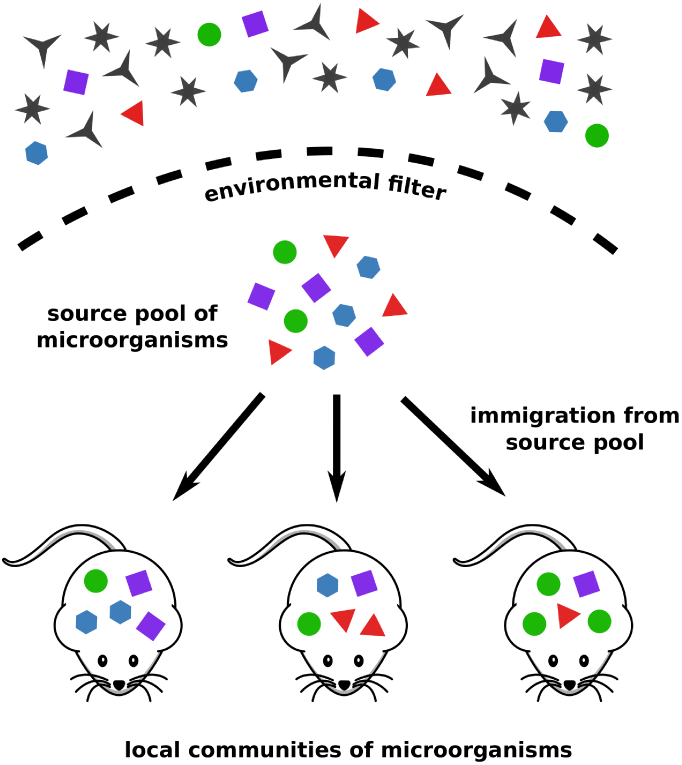
Sketch of the setup of the neutral model. Only a subset of the available microorganisms can pass through the environmental filter, forming a common source pool for all hosts. This selective step is not described by the neutral model. The microbiota of individual hosts form local communities and the microbial intra–host population dynamics are neutral. Local diversity is maintained by immigration from the source pool.

Despite these controversies and with the growing availability of microbial community data, the neutral theory has been adapted and applied to the microbial world [20, 21, 22, 23, 24]. In particular, the local community vs. metacommunity structure of neutral theory naturally extends to host-associated microbial communities. Here, the hosts are viewed as ecosystems and their microbiota are treated as local communities [25, 11]. This has led to several recent studies addressing the question whether the microbiota conform to the patterns predicted by the neutral theory. The rank-abundance patterns of the microbiota from three domesticated vertebrates for example were found to be largely consistent with the neutral expectation [26]. A good fit of the neutral model was also found for the microbiota of young individuals of the zebrafish *Danio rerio* [27]. Interestingly, in this study neutrality decreased as hosts aged, indicating an increasing influence of selective processes over developmental time, potentially linked to the activation of a fully functioning adaptive immune response. The human microbiota were found to be predominantly non-neutral across various body sites [28], which is thought to be the result of a phase transition from a dispersal-dominated neutral regime to a selection-dominated within-host regime [29]. In this study the few neutral communities were mostly associated with the urological tract and the skin, while in another study the composition of the skin microbiota of healthy human subjects in large Chinese cities were found to be better explained by non-neutral processes [30]. Contrasting results can also be obtained depending on the state of the host, with the healthy human lung microbiota being largely consistent with a neutral model, while microbes recovered from diseased lungs diverged from neutrality [31]. Neutral assembly processes in invertebrate hosts are understudied, but a recent study shows the microbial communities associated with the fruit fly *Drosophila melanogaster* to be consistent with the predictions of a neutral model [32]. While these studies have drawn awareness to the potentially important contribution of neutral processes in shaping the microbiota, the use of several different neutral models and focus on one or a few, mostly vertebrate host organisms, makes it hard to draw more general conclusions.

Here, we aim to overcome these limitations by consistently applying a neutral model to a variety of distinct host systems, including, but not limited to, a wide range of invertebrate hosts. We included host species from a range of eukaryotic multicellular organisms with different life styles, ranging from early branching groups such as sponges to the house mouse with its fully developed adaptive immune system. We used a general modelling approach and combined it with a consistent simulation framework, in order to achieve an unbiased comparison of the distinct metaorganisms. Based on the neutral null hypothesis, we identified members of the microbiota which consistently deviate from the neutral expectation, possibly indicating differential selection of these particular microbes. Additionally we hypothesize that some instances of observed non-neutrality may be due to transient community stages, where the microbiota have not yet reached its long-term equilibrium composition.

## Results

We employed the neutral model presented by Sloan et al. [20], which is particularly suited for large-sized microbial populations and has been used before to assess microbiota neutrality [27, 32]. It describes the stochastic population dynamics within a local community, corresponding to the microbiota of a specific host. To maintain local diversity, which would otherwise reduce to a single species through ecological drift, immigration from a fixed source community occurs with rate *m*. This source community is not equivalent to the environmental pool of microorganisms, but rather it can be interpreted as the collection of all microbial species that can pass through the environmental filter of the host (Fig 1). Thus, the neutral model does not make any assumptions about the selective constraints that apply before and during colonization of the host and in particular, it is not concerned with any host traits that may restrict the range of colonizing microbes [33].

This model allows to derive an expression for the expected long-term stationary community composition. Specifically, it predicts the relationship between the mean relative abundance of a taxon across all local communities, i.e. the metacommunity, and the probability of actually observing this taxon in any single community. It has been shown that this relationship is determined by a beta distribution ([20] and *Materials and Methods*). The only free parameter of this model is the immigration rate *m*, which can be calibrated by a nonlinear fit. The goodness of the fit then indicates how well the prediction of the neutral model compares to the empirical data. More specifically, we used the coefficient of determination *R*^2^ as a quantitative measure of how consistent the data is with the neutral model. See the *Material and Methods* for details of the neutral model, simulations and the fitting procedure.

### Neutrality of the microbiota

We fitted the theoretical neutral expectation to the published microbiota compositions of eight different host species across four phyla and, for comparison, three environmental microbial communities (see the *Materials and Methods* and S1 Table for an overview and references).

With the species *Sarcotragus fasciculatus, Ircinia oros* and *Carteriospongia foliascens*, our study included examples from the oldest extant sister group to all other animals, the sponges (Porifera) [34]. Despite their filter-feeding life-style, sponges harbor a very diverse and highly specific microbiota [35], which mediates the functional role of the sponges in the ecosystem [36]. This is complemented by the jellyfish *Aurelia aurita*, the starlet sea anemone *Nematostella vectensis*, and the fresh-water polyp *Hydra vulgaris*. These hosts are examples from another early branching phylum, the Cnidaria, which possess a basic innate immune system thought to play an important role in controlling the interaction with the associated microbial community [37]. We also included the nematode *Caenorhabditis elegans* as one of the best–studied multi-cellular organisms, whose microbiota has only more recently come into the focus [38, 39, 40, 41]. Finally, the house mouse *Mus musculus* with its potent adaptive immune system [42] offers a well–studied intestinal microbiota [43, 44]. Our datasets included lab-reared animals as well samples from several natural populations from various locations, representing a broad range of environmental and microbial contexts. For some host species samples from both lab and natural populations were available, which we analyzed separately to assess the potential impact of heterogeneous natural habitats vs. homogeneous, constant lab conditions, on neutrality. See the *Materials and Methods* for a summary of the context of the individual datasets.

In all cases, community composition was determined by the relative abundances of operational taxonomic units (OTUs), obtained by standard high-throughput 16S rRNA sequencing techniques [45]. To ensure equal sample sizes, all OTU abundance tables were rarefied to the same read depth (1000 reads per sample). We analyzed the effect of the rarefaction on the neutral fit for datasets where more reads were available and found it to be of little relevance for most communities (S6 Fig).

A comparison of the theoretical and observed relationship between mean relative abundances and occurrence frequencies of OTUs for three illustrative examples is shown in Fig 2 (see S1 Fig for examples of the neutral fit for all datasets). The neutral model generally predicts a specific monotic increase of the occurence frequency of an OTU with an increasing mean relative abundance of this OTU across all samples. This is a quantitative reflection of the null expectation that more abundant OTUs should also be found in more samples. Consequentely, OTUs that are found below the prediction in the lower right region of the graph are found in fewer samples than expected by their mean abundance across all samples. Conversely, OTUs that lie above the neutral expectation are found more often than expected.

**Figure 2.**
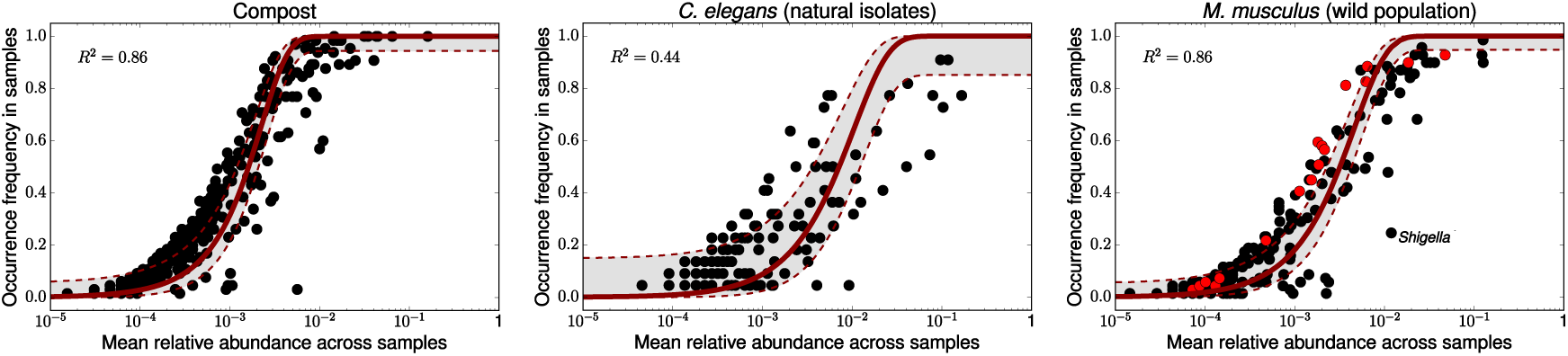
The relationship between mean relative abundance of taxa across all samples and the frequency with which they are detected in individual samples for three representative datasets. Each dot represents a taxon and the solid line is the best-fitting neutral community expectation. The dashed lines and grey area depict the 95% confidence bands. The left panel shows a microbial community sampled from a compost as an example of an abiotic environment. The centre panel shows data for the natural microbiota of *C. elegans* nematode populations sampled from the same compost. The right panel shows the microbiota of a wild *M. musculus* mouse population. Here, the red dots indicate members of the microbial family Ruminococcaceae, which are predominantly overrepresented compared to the neutral expectation. A genus that was found in much fewer mice than expected by chance is *Shigella*, being present in less than a quarter of the hosts despite a neutral expectation of nearly 100% prevalence.

A summary of the goodness of fit of the neutral model to the microbial communities from all datasets is shown in Fig 3. Higher *R*^2^ values generally indicate a closer match between the neutral expectation and the data. To aid the interpretation and comparison across datasets, we additionally contrasted every dataset with a corresponding simulated, completely neutral community that had the same sample size, microbial diversity and estimated immigration parameter *m* (see Material and Methods for details of the simulation).

**Figure 3.**
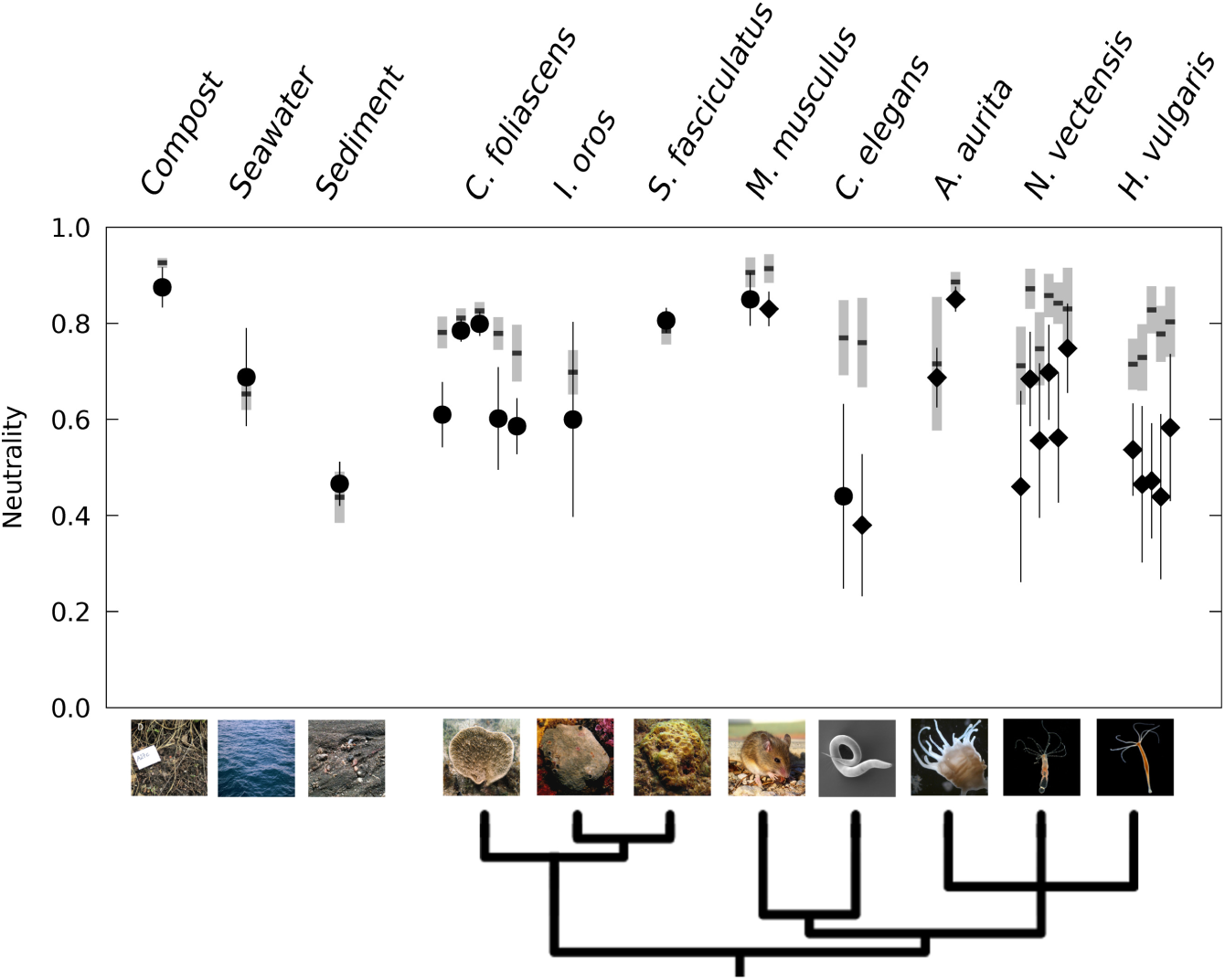
Goodness of fit of the neutral expectation to a range of host-associated and environmental communities. Circles denote natural populations and diamonds denote laboratory populations, error bars indicate 95% bootstrap confidence intervals. The grey bars indicate the 95% bootstrap confidence intervals around the expected goodness of fit (horizontal black line) to a neutral simulation, providing a comparison to a completely neutral community with the same sample size and diversity. The data for *C. foliascens* is from several different natural populations, while the data for *H. vulgaris* and *N. vectensis* is from different time points. Spread of points along the x-axis is added to increase visibility. Phylogeny generated with *phyloT* based on NCBI taxonomy. Sponge photographs courtesy of Susanna López-Legentil (UNC Wilmington) and Mari-Carmen Pineda (AIMS).

This allows a comparison of the deviation between the empirically obtained measure of neutrality and the expected values from a known neutral community for each dataset. This is crucial, since depending on the sample size and microbial diversity, even a completely neutral community will not yield a perfect fit, even when there is sufficient time to reach an equilibrium (Fig 4).

**Figure 4.**
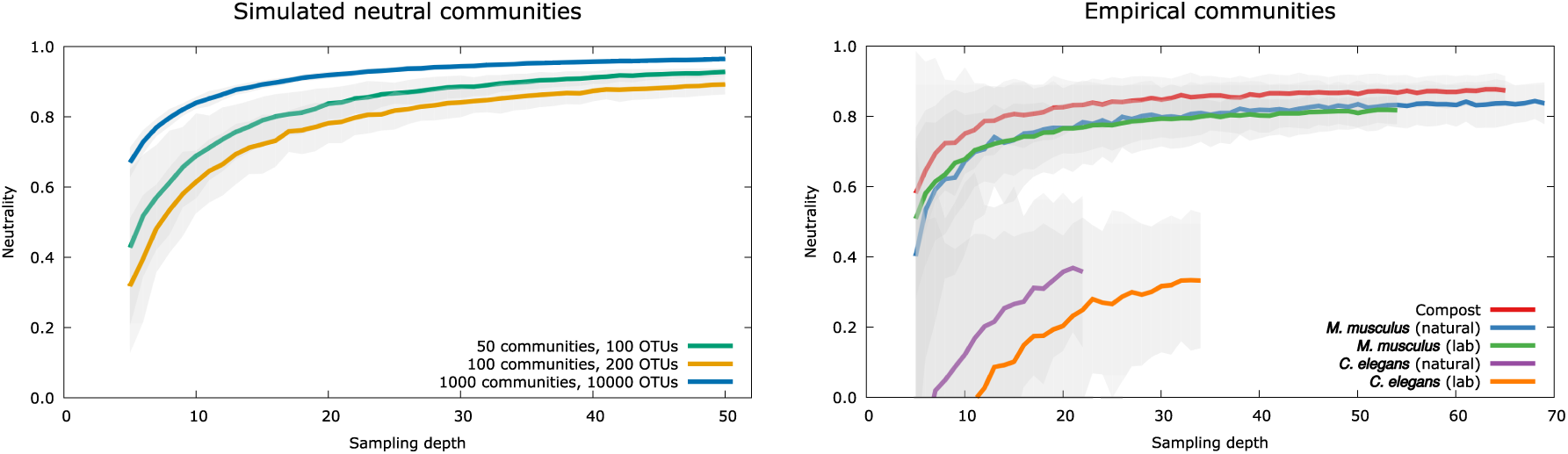
Impact of sample number on the goodness of fit of the neutral expectation. For each sampling depth, communities where resampled 100 times with replacement at a rarefaction level of 1000 reads/sample. The left panel shows the results for simulated neutral communities. The immigration parameter was fixed at *m* = 0.05 and neutral dynamics were simulated for 10^6^ timesteps in all cases. Generally goodness of fit remained high over a large range of sample sizes, reflecting the underlying completely neutral community dynamics. Note, that goodness of fit did not reach 1.0 in finite and relatively small metacommunities, even if all communities were sampled. The right panel shows the effect of subsampling for several host-associated microbial communities. As for the simulated data, observed levels of neutrality were relatively robust, only dropping off at very low sample sizes. The *C. elegans* datasets showed a very poor fit at very low sampling depths, but began to level off well below the neutrality values of the other datasets. Shaded areas indicate 95% bootstrap confidence intervals.

Turning first to the environmental microbial communities, we generally observed a high consistency with the neutral expectation. The majority of microbial taxa found in compost samples for example closely followed the neutral prediction (Fig 2), as reflected by a high goodness of fit measure (*R*^2^ = 0.86). While the goodness of fits to the microbial communities in seawater and marine sediment are lower, they lie in the range expected from the respective neutral simulations. The sediment dataset showed a marked effect of rarefaction, with the goodness of fit increasing to the same level as the seawater community when more reads are included (S6 Fig). It is also the dataset with the highest microbial diversity (*>* 3000 OTUs) and a relatively low sample size, which may indicate a greater influence of rarefaction on rare taxa. The overall good consistency of the environmental communities with the neutral expectation aligns with previous studies showing that random population dynamics and immigration play a major role in shaping microbial communities in abiotic environments [21, 23, 24].

Biological hosts with their potential for active selection of specific taxa, could on the other hand be expected to have their resident microbial communities under tighter control, reducing the relative importance of stochastic processes after colonization, which would be reflected in a worse fit of the neutral expectation. The nematode *C. elegans* for example naturally consumes bacteria as its food source and possesses an innate immune system as a defence against the threat of ingested pathogenic microbes [46]. This suggests that these worms have some control over their microbiota and indeed we found a substantial mismatch between the neutral expectation and the worm microbiota (Fig 2), resulting in a low goodness of fit (*R*^2^ = 0.44). We also found a similar level of neutrality for the microbiota of laboratory worm populations (Fig 3), indicating a similar influence of neutral processes under more controlled laboratory conditions. For both the natural and lab-enriched worms the goodness of fit was well below the levels expected from the corresponding neutrally simulated communities (Fig 3). It is noteworthy that these specific worms were isolated from the same compost where the environmental microbial communities showed a very good alignment with the neutral expectation (Fig 2, 3).

Vertebrate hosts with their more sophisticated and complex immune system are expected to diverge even further from the neutral expectation. However, intriguingly, the neutral model showed a very good fit to the microbiota of a wild *M. musculus* mouse population (Fig 2). The best fit was in fact on par with the fit to the environmental microbes isolated from compost (*R*^2^ = 0.86) and the deviation from the simulated neutral community was minimal (Fig 3). The high level of neutrality in the natural mouse population was corroborated by a laboratory population of *M. musculus* (Fig 3).

We found similar high levels of neutrality for sponges, which as aquatic filter feeders are exposed to the full range of marine microbes. Like nematodes, they have a basic immune system [47], and especially for *S. fasciculatus* and some populations of *C. foliascens*, the data was in very good agreement with the neutral expectation (Fig 3, S1 Fig). In contrast, the results for the three aquatic polyps were less consistent. The two laboratory populations of *A. aurita* were in very good agreement with the neutral prediction, while samples taken at different developmental stages of *H. vulgaris* and *N. vectensis* generally showed a larger deviation from full neutrality (Fig 3).

The estimated immigration parameter *m*, which can be interpreted as a proxy for inter-host transmission and similarity of the individual microbial communities, showed considerable variation across the datasets (S2 Fig). Overall, communities from aquatic samples showed higher best-fit values of *m* than terrestrial samples. This potentially indicates an increased between-sample dispersal and a more homogeneous microbial distribution in aquatic settings, at least on the relevant spatial scales. This seems especially plausible for the filter-feeding sponges sampled from the same location. In contrast, we obtained relatively low estimates for the immigration parameter for the mice and nematodes, in both the natural and the laboratory populations. For nematodes in particular, a more patchy distribution of microbes on the spatial microscale, together with stochastic founder effects during colonization [48], may play a role in creating more heterogeneous microbiota compositions. In combination with less exchange of microbes in this terrestrial organism with low dispersal ability, this can contribute to low values of *m*.

Note, that we did not find a correlation between immigration rate *m*, sample size or microbiota diversity and the level of neutrality (S5 Fig). We additionally analyzed the effect of varying sample sizes by subsampling all datasets with more than 20 samples, since the number of individual samples has the potential to influence the composition of the source community of available microbial species. We found the observed levels of neutrality to be robust against subsampling, unless sample sizes are very small (Fig 4). In all cases, neutrality consistently decreased with decreasing sample sizes, potentially reflecting a decreasing overlap in the local microbial communities. We performed this analysis for both the simulated and the empirical datasets, which both showed a very similar impact of subsampling.

Comparing the neutral expectation with data from a wide range of environments and host species showed that a good fit of the neutral model is in fact the norm rather than the exception (Fig 3). In particular, there was no substantial difference in observed neutrality between microbial communities living in environmental samples, such as compost and seawater, and the microbiota from several animal host species. We thus conclude that the observed patterns of microbiota composition in hosts as different as mice and marine sponges can in many cases already be obtained with a neutral null model.

### Identifying non-neutral taxa

Our analysis showed that for all host species, regardless of the specific goodness of fit of the neutral model, there are microbial taxa that diverged substantially from the neutral expectation. Since an essential ingredient of neutral theory is the assumption of selective equivalence, such a divergence from neutrality may signify taxa that are subject to differential (e.g. host-mediated) selection. Thus, identifying specific non-neutral members in the bulk of the microbiota is of great practical importance. In particular, OTUs that are observed more often than expected by chance are commonly seen as good candidates for beneficial symbiotic microbes. In general, however, it is not straightforward to decide whether a microbial taxon is under positive or negative selection within a specific host. A taxon could for example be observed more often than expected because its is indeed essential for the host even at low abundances, and is thus positively selected for in the community. Alternatively, such over-representation may point to a microbe which is simply a very good colonizer, i.e. it can easily disperse and pass through a host’s environmental filter, but within the host it is subject to negative selection to prevent it from reaching high abundances. Here, the practical value of the neutral model lies in its ability to identify microbes that appear to interact in a specific, non-neutral form with the host and are thus worthy of further investigation, taking into consideration ecological and functional information.

In the following, we classified all OTUs outside of the 95% confidence bands around the neutral prediction as not consistent with the neutral expectation. We found that the distribution of non-neutral OTUs around the neutral prediction is roughly symmetrical in all cases, and no microbial order deviated systematically above or below the neutral expectation (S3 Fig).

To further distinguish random deviations from neutrality and actual non-neutral candidates, we also applied a more conservative definition of non-neutrality. For this, a particular OTU was classified as non-neutral only when it consistently diverged from the neutral expectation in the same direction in independent host populations. Applying this definition to the natural and laboratory populations of *C. elegans* and *M. musculus* yielded a reduced subset of microbial taxa, which lay above or below the neutral prediction in both populations.

This analysis revealed that of the *C. elegans* microbiota, only the genus *Ochrobactrum* is consistently under-represented (S2 Table). It is found in only ca. 40% of the natural isolates and 70% of the laboratory worms, despite its relatively high mean abundance across all worms and a neutral expectation of 100% prevalence. This underrepresentation of *Ochrobactrum* is intriguing, as it had previously been identified to be enriched in worms, to favor growth of worm populations, and to be able to persist in the nematode’s gut even under starvation conditions [39]. However, this observation is consistent with a transient community state (see below), where *Ochrobactrum* has not yet colonized all worms, despite being very successful once it has entered the worm.

In the *M. musculus* microbiota of natural and laboratory populations, over a third of the over-represented taxa were from the bacterial family *Ruminococcaceae*. Members of this family were never consistently found below the neutral expectation across hosts (Fig 2 and S3 Table). This family includes several physiologically relevant genera involved in the utilization of plant polysaccharides such as starch and cellulose [49], and high-fat diets low in plant-derived materials have been found to decrease the proportion of the *Ruminococcaceae* in mouse guts [50]. A notable microbe which was found in significantly fewer mice than expected from the neutral model is *Shigella* (Fig 2). Invasive *Shigella* can cause intestinal inflammations in humans, but mice mount an effective defense against it, thereby preventing acute infections [51]. This illustrates how potentially pathogenic microbes, which may be actively selected against by the host, can be found diverging from the neutral expectation. That we were able to consistently identify a bacterial family that had previously been linked to important metabolic functions as over-represented and potential pathogens as underrepresented, shows the utility of the neutral model as a null hypothesis and a tool of finding meaningful patterns in the vast complexity of the microbiota.

Another commonly applied approach to tackle this complexity consists of the identification of OTUs that are shared among hosts, typically denoted as the core microbiota. This usually requires setting a specific occurence threshold for the taxa found across hosts in similar habitats [52]. Applying this persistence-based definition to our data shows that a substantial portion of the core taxa, e.g. those with an occurence frequency above 90%, fall within the neutral expectation (Fig 2, S1 Fig). This implies that many taxa within the abundant and prevalent core microbiota may actually not occur more often than would be expected by chance. Taxa such as the *Ruminococcaceae* in mice on the other hand, which were below typical cut-off prevalences but occurred more often than expected by chance, emerge as interesting candidates beyond the usual core taxa. Deviations from the neutral expectation can thus complement and enrich insights gained from the widely used concept of the core microbiome.

### Change of neutrality over time

As we have discussed in the previous section, the most commonly assumed cause of deviations from neutrality is selection by the host, for example through its immune system. In the context of our study this would mean that after colonization, host-mediated selection may have altered the composition of the microbial community, resulting in a non-neutral community structure. Interestingly, we found a particularly strong deviation from neutrality for the nematode host, which possesses a rather simple immune system [53]. In contrast, other hosts with more complex immune systems, for which strong selection could have been presumed, show microbiomes close to neutral expectations. This suggests a role of additional factors beyond the immune system driving deviations from the neutral expectation.

A crucial aspect of the neutral prediction is that it is based on the expected *long-term* stationary distribution of the microbial community. If the community is in a transient state, however, it can appear non-neutral even if it is the product of a purely neutral process. This effect is illustrated in Figure 5, showing a simulation of the population dynamics of 50 neutral communities (*Material and Methods*). For every community, only a few random colonisers were picked from the source community, which yields random, but non-neutral, initial communities. We then compared the neutral expectation to the in-silico communities over time during the simulation, showing that they become “neutralized” through random population dynamics and immigration until the community composition reaches its long-term equilibrium, or the host dies.

**Figure 5.**
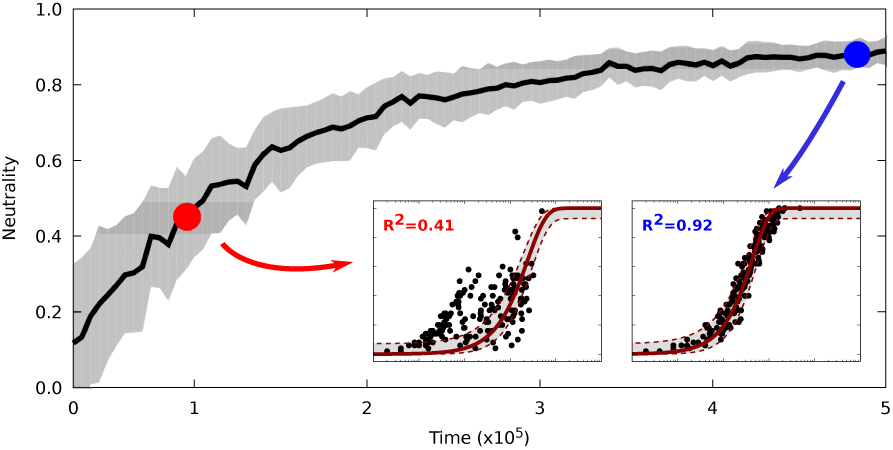
Goodness of fit of the neutral long-term expectation to neutrally simulated communities over time. The two insets show the best fits of the neutral model to snapshots of the communities at an early timepoint (indicated by the red dot) and a late time-point (indicated by the blue dot). Scale and meaning of the axes for the insets are the same as in Fig 2. Each community was initialised with a few random colonizers, which yielded non-neutral initial communities that neutralized over time through a stochastic death-birth process and immigration. The shaded area indicates the 95% bootstrap confidence intervals.

Thus, if community assembly is governed by purely neutral processes, we would expect the fit of the neutral model generally to become better over time. Decreasing neutrality, which has been reported for the zebrafish *Danio rerio* [27], would on the other hand be a good indicator of selective processes. A similar increase in the relative importance of selective processes over stochastic effects has been described for the ecological succession of microbial communities in salt marshes [54]. However, while the microbiota from the *Hydra* and *Nematostella* populations show clear developmental patterns [55], we did not find a clear temporal trend in the deviation from the neutral model (S4 Fig).

While we could not conclusively disentangle transient effects in the interplay between ecological drift and selective processes, in principle it is possible that a poor fit of the neutral expectation may be due to a transient non-equilibrium state of the community, rather than actual non-neutral processes. This effect would be especially pronounced for short-lived host species such as *C. elegans*, where initial colonization can lead to communities that appear non-neutral even for ecologically equivalent (i.e. neutral) colonizers [48]. The microbiota of such hosts may never effectively converge to the neutral long-term expectation within the lifetime of the host, even if they are dominated by stochastic dynamics and immigration. For longer–lived species, such as sponges and mice, we would expect these transient effects to play a smaller role.

### Comparison with a random sampling model

The observed high levels of neutrality do not necessarily imply that the microbiota are simply a random collection of microbes. A good fit of the expected distribution indicates that the simple neutral model is sufficient to describe the relationship between abundance of a species and its occurrence frequency. But this can not rule out other processes, niche-related or stochastic, that lead to the same community structure [56].

A very simple alternative model is obtained by assembling a local community by drawing randomly from the fixed source community, thus neglecting stochastic reproduction events and dispersal. In this case, the probability of observing a specific species in a local community is determined by a binomial distribution. Comparing the best fits of the neutral expectation and of a binomial distribution using the Aikake information criterion (AIC) indicated that the neutral model fits the data better in all cases (S1 Table). This indicates that the microbiota are not merely random samples of the microbes present in the metacommunity.

In a more general context, ecological communities such as microbiota are examples of component systems, where each specific instance of the system (e.g. a specific microbial community) is comprised of components drawn from a shared set of basic building blocks (e.g. the source pool in our scenario). Such systems have been shown to exhibit some general invariant properties, and deviations from these properties may reveal functional constraints imposed on the shared components [57].

## Discussion

We set out to quantify how consistent the microbiota of a range of different host organisms are with the expectation from a neutral model. We found that the structures of the considered host-associated microbial communities are often in surprisingly good agreement with the neutral expectation. Taking into account a more complete simulation analysis of the neutral model further suggested that transient stages of the microbiota may contribute to the divergence from neutrality observed for some host species.

Even a high consistency with the neutral expectation does not rule out the presence of selective abiotic or biotic processes, as non-neutral models can yield predictions consistent with neutral theory under certain conditions [56]. Moreover, biological hosts will select for a subset of the environmentally “available” microbes, and yet the resulting microbiota may still look mostly neutral. In fact, comparing the microbes present in seawater to the species present in individual sponges reveals that there is almost no overlap (S7 Fig). This indicates that sponges are a highly selective environment, imposing strong constraints on potential colonizers, and yet after colonization their microbiota are very consistent with the neutral expectation. A similar selective filter is found for *C. elegans*, where only a fraction of the microbes present in the environment successfully colonize the worms (S7 Fig). This suggests that there are host traits that act as environmental filters [33] and they often will do so in a predictable, i.e. deterministic way. In this sense the microbiological tenet that “*everything is everywhere*, but, *the environment selects*” [58] also applies when the environment happens to be another biological organism. However, our results highlight that the processes that are at play *after* the environmental filter has been passed can be highly consistent with the neutral model. This apparent contradiction between selective environments and neutral processes acting on the allowed set of species may also have contributed to some of the contrasting results found in previous studies [59].

This filtering of potential colonizers is not unique to biological hosts, but what sets biological hosts apart from other habitats is the potential of the hosts’ filtering mechanisms to evolve. An example for such an evolvable mechanism is the production of antimicrobial peptides (AMPs) by several *Hydra* species [60]. Here, each *Hydra* species was shown to posses a specific AMP expression profile, selecting for a species-specific set of bacteria by inhibiting colonization of foreign microbes. Another recent study showed how host-derived proteins from the jellyfish *A. aurita* interfere with bacterial quorum sensing to modulate host colonization [61]. While these examples do not constitute proof of adaptive coevolution between the host and its microbiota, they exemplify the possible specific selection capabilities of the host. Moreover, our results then highlight that microbiota composition is the outcome of several intertwined processes. For example, selective filtering at the colonization stage may be controlled by the host, or by microbe characteristics, and is then presumably followed by largely neutral, stochastic processes. It is further possible that selection at either the colonization stage or community assembly thereafter only affects a subset of microbes, while the remaining microbes follow neutral dynamics. Another factor to consider is that taxonomic and functional microbial community composition may be shaped by largely independent processes [62, 63]. Given the known examples of functional redundancy present e.g. in the human gut microbiota [64], viewing communities from the perspective of functional (e.g. metabolic) categories can provide an alternative picture to data based on 16S rRNA gene based taxonomic profiling [65]. These intertwined processes need to be better understood and taken into account for an enhanced understanding of microbiota assembly.

Our simulation analysis demonstrates that the community needs time to reach the dynamic neutral equilibrium (Fig 5) through ecological drift. This emphasizes the need for temporal data to further investigate the role of transient community stages and drift in microbial communities. Such data is usually not available for host-associated microbial communities, and our study highlights the need to be aware of this potentially confounding factor when applying neutral models to cross-sectional community snapshots.

The neutral assumption that species do not interact is an anathema to many ecologists, and for the microbiota in particular it is assumed that the functioning of the community is enhanced by cooperative interactions. Recent results however suggest that interactions within the mouse gut microbiota are indeed predominantly competitive and very weak [66]. Together with our finding of the microbiota often being consistent with neutral expectations for several different host species, this lends support to taking the neutral null hypothesis as a key component of our understanding of host-microbe interactions. We would like to emphasize that even a good consistency with the neutral model does not imply that the microbiota are functionless or that hosts are non-selective – the *Hydra* system actually contains taxa that confer a selective advantage, but which appear as neutral in our present analysis [67]. Neutral theory in fact makes no claims about microbiota function, rather it assumes that the different community compositions that arise after the environmental filter of the host has been passed are, in the words of the original formulation of the neutral theory, *selectively nearly equivalent, that is, they can do the job equally well in terms of survival and reproduction* [68] of the individual host.

Awareness of the potentially significant role of neutrality in shaping the microbiota is crucial to avoid being led astray by randomness in view of the incredible complexity of host-microbiota symbioses and to be able to uncover general principles of microbiome assembly.

## Methods

### Summary of datasets and data availability

The collection of microbial abundance datasets for the environmental and host-associated communities (i.e. OTU tables) was assembled as part of the Collaborative Research Centre 1182 “Origin and Function of Metaorganisms” and is available from the authors on request. The raw sequencing data has been published previously and is available online, see S1 Table for the corresponding references. In the following we give a short summary of the context of the respective datasets, for details we refer to the corresponding references.

#### Sponges [35]

*S. fasciculatus* and *I. oros* samples were collected at the same location on the Spanish coast on different dates. *C. foliascens* samples were collected at different locations along the Australian coast on different dates.

#### C. elegans [39]

The natural isolates of single *C. elegans* worms were collected on different dates at the same location in Northern Germany.For the lab enriched populations of *C. elegans*, single worms isolated from the same location were transfered to agar plates seeded with *E. coli* as food. On these plates the worms were allowed to reproduce via selfing and were maintained for 2-3 weeks before sampling the resulting populations.

#### Mice [44]

For the natural mice, 69 mice were captured from 34 unique locations, with a maximum of 3 mice per location, in the hybrid zone of *M. musculus musculus* and *M. musculus domesticus* in Bavaria, Germany, during May and June 2011.

The lab dataset includes 40 F2 hybrids between *M. musculus domesticus* and *M. musculus musculus* standard lab strains, and additionally seven each of the *domesticus* and *musculus* parental mice.The parental strains were kept under conventional conditions for two generations before setting up experimental crosses.

#### *N. vectensis* [55]

Animals were sampled from 15 independent laboratory cultures under different, but fixed environmental conditions (temperature and salinity) at specific time points.

#### *H. vulgaris* [69]

Animals were sampled at several time points from independent, clonal lab cultures with identical and constant environmental conditions.

#### *A. aurita* [61]

Animals were sampled from 4 independent populations, one control population and 3 treatment populations.The treatment (QQ) populations were each treated with a different host-derived quorum-quenching protein.

#### Environmental samples

Compost samples were collected in the context of the *C. elegans* study [39], following the same protocols and at the same locations.Marine seawater and sediment samples were collected in the context of the sponge microbiota study [35].

### The neutral model

Here, we summarize the model presented in [20], considering only the purely neutral special case with equivalent growth rates for all species. A local community contains a fixed number of *N* individuals, of which *N*_*i*_ are of species *i*. Individuals die with a constant death rate *δ* and a dead individual is replaced by reproduction of a local individual (with probability 1 *− m*), or by immigration from a fixed source community (with probability *m*). Thus the total population size *N* remains constant. The relative abundance of a species in the source community is denoted by *p*_*i*_. With this, the probabilities that the number of individuals of species *i* increases by one, decreases by one or stays the same are given by

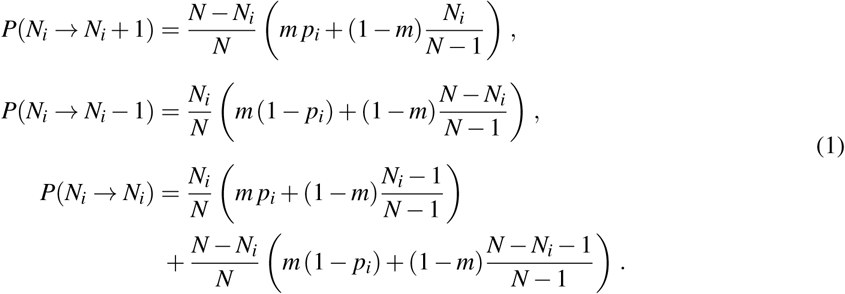

These transition probabilities correspond to Hubbell’s original neutral model, but the following continuous approximation derived by Sloan et al. [20] can be efficiently applied to very large population sizes and allows for a relatively simple analytical solution. In particular, this allows for a calibration with the high-throughput 16S rRNA sequencing data obtained from microbial communities.

If *N* is large enough, such as in microbial communities, we can assume the relative abundance *x*_*i*_ = *N*_*i*_/*N* of species *i* to be continuous. This leads to a Fokker-Planck equation for the probability density function *ϕ*_*i*_ (*x*_*i*_, *t*) of *x*_*i*_,

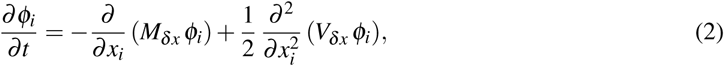

where *M*_*δ x*_ and *V*_*δ x*_ are the expected rates of change in relative abundance and variability, respectively. Assuming the time intervals between individual death-birth events are short, they can be approximated by

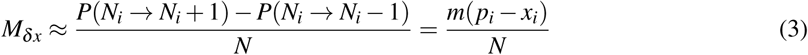

and

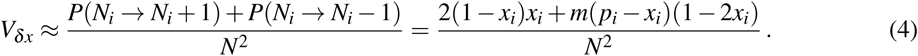

Equation (2) describes the neutral population dynamics of a local microbial community. This is not in general amenable to analytical treatment, but one can approximate the long-term equilibrium solution of this equation. Namely, the so-called potential solution for *∂ϕ_i_/∂t* = 0 arising from 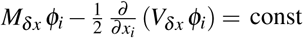., is given by the beta distribution

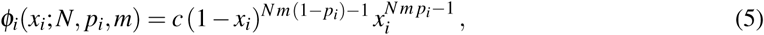

with *c* = Γ(*N m*)*/* [Γ(*N m* (1*- p_i_*)) Γ(*N m p_i_*)]. This can be connected to empirical observations. For a given detection threshold *d*, the probability of actually observing a species in a local community is given by the truncated cumulative probability density function

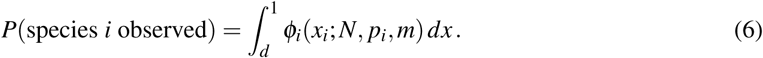

Here, *N* is given by the number of reads per sample and the relative abundance *p*_*i*_ of the focal species in the source community can be approximated by the mean relative abundance of the species across all samples. The detection threshold is set to *d* = 1*/N*. The probability of immigration *m* is thus the only free parameter and can be used to fit the predicted long-term distribution to the observed occurence frequencies of the microbiota.

### Fitting procedure

The fitting process applied to all datasets, empirical and simulated, works as follows. First, the OTU abundance table was rarefied to the desired read depth (*N* =1000 reads per sample for the main results, which was determined by the lowest available read depth from all datasets). The expected observation probability (6) and a binomial model were then fitted to the observed mean relative OTU abundances *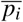* and occurence frequencies *f*_*i*_ obtained from this rarefied table using non-linear least squares minimization with the *lmfit* package (pypi.org/project/lmfit) [70]. As a measure of the goodness of fit we then calculated the standard coefficient of determination using the ratio of the sum of squared residuals and the total sum of squares:

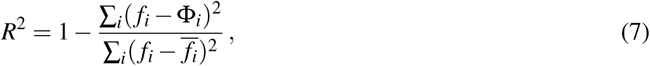

where Φ*_i_* is the expected ocurrence frequency obtained from the best-fit neutral prediction. Additionally, for comparison of the neutral and binomial model, the mean Aikake information criterion (AIC) was calculated. In all cases, 95% bootstrap confidence intervals were obtained by resampling the hosts 100 times with replacement and performing the described fitting procedure on each resampled set of hosts.

### Neutral model simulations

Simulated neutral communities were generated following the stochastic death-birth-immigration process described above and in [20]. Communities with a prescribed number of samples/communities, community size, source pool diversity and immigration parameter *m* were started from random initial conditions and simulated for 10^6^ time steps to let them reach dynamic equilibrium. The community snapshots obtained at the end of each simulation were treated with the same fitting procedure as the empirical datasets.

### Code availibility

The Python code for fitting and simulating the neutral model is available at http://github.com/misieber/neufit.

## Neutrality in the Metaorganism - Supporting Information

**Table S1.** Overview of the datasets included in this study, their source (natural or laboratory communities), number of samples, microbial diversity and a summary of the results of the model fits. Data is for rarefaction level of 1000 reads per sample. Reference is to the original study containing detailed information on sample preparation and raw sequencing data.

| Samples | Source | #samples | #OTUs | best-fit $m$ | $R^2$<br>neutral | AIC<br>neutral | binomial | Reference |
| --- | --- | --- | --- | --- | --- | --- | --- | --- |
| *Caenorhabditis elegans* |  |  |  |  |  |  |  | [39] |
| natural samples | natural | 22 | 193 | 0.03 | 0.44 | -673.2 | -452.5 |  |
| lab samples | lab | 34 | 106 | 0.007 | 0.38 | -330.0 | -208.2 |  |
| *Ircinia oros* |  |  |  |  |  |  |  |  |
| Spain, Barcelona | natural | 11 | 1121 | 0.41 | 0.6 | -5391.47 | -4966.26 | [35] |
| *Sarcotragus fasciculatus* |  |  |  |  |  |  |  |  |
| Spain, Barcelona | natural | 12 | 735 | 0.83 | 0.81 | -3552.82 | -3165.09 | [35] |
| *Carteriospongia foliascens* |  |  |  |  |  |  |  | [35] |
| Australia, Davies Reef | natural | 15 | 939 | 0.36 | 0.6 | -5435.40 | -5007.39 |  |
| Australia, Fantome Island | natural | 14 | 728 | 0.81 | 0.79 | -3740.45 | -3336.08 |  |
| Australia, Orpheus Island | natural | 15 | 750 | 0.86 | 0.8 | -3987.01 | -3539.84 |  |
| Australia, Green Island | natural | 13 | 774 | 0.47 | 0.59 | -3846.0 | -3542.79 |  |
| Australia, Torres Strait | natural | 7 | 336 | 0.78 | 0.58 | -1410.75 | -1244.0 |  |
| *Mus musculus* |  |  |  |  |  |  |  | [44] |
| natural samples | natural | 69 | 281 | 0.11 | 0.85 | -1217.97 | -998.15 |  |
| lab samples | lab | 54 | 136 | 0.18 | 0.84 | -536.30 | -452.48 |  |
| *Nematostella vectensis* |  |  |  |  |  |  |  | [55] |
| 1 day post fertilization (dpf) | lab | 6 | 226 | 0.23 | 0.52 | -738.68 | -655.06 |  |
| 4 dpf | lab | 15 | 149 | 0.33 | 0.67 | -521.62 | -479.29 |  |
| 40 dpf | lab | 12 | 155 | 0.09 | 0.59 | -502.39 | -413.96 |  |
| 123 dpf | lab | 12 | 195 | 0.39 | 0.69 | -678.72 | -623.49 |  |
| 385 dpf | lab | 20 | 225 | 0.17 | 0.58 | -766.93 | -679.74 |  |
| 401 dpf | lab | 8 | 120 | 0.7 | 0.74 | -446.34 | -402.59 |  |
| *Hydra vulgaris* |  |  |  |  |  |  |  | [69] |
| 0.5 weeks after hatching (wah) | lab | 8 | 699 | 0.6 | 0.5 | -2527.91 | -2289.35 |  |
| 2.5 wah | lab | 8 | 248 | 0.28 | 0.44 | -820.92 | -756.19 |  |
| 5 wah | lab | 8 | 242 | 0.53 | 0.43 | -1129.85 | -1041.28 |  |
| 9 wah | lab | 8 | 257 | 0.5 | 0.44 | -833.46 | -765.88 |  |
| 15 wah | lab | 8 | 140 | 0.45 | 0.59 | -466.20 | -432.55 |  |
| *Aurelia aurita* |  |  |  |  |  |  |  | [61] |
| control | lab | 5 | 163 | 0.91 | 0.69 | -624.96 | -538.45 |  |
| quorum quenching (QQ) | lab | 18 | 391 | 0.62 | 0.85 | -1716.09 | -1532.6 |  |
| Environment |  |  |  |  |  |  |  |  |
| Compost | natural | 65 | 587 | 0.48 | 0.86 | -2690.26 | -2491.77 | [39] |
| Seawater | natural | 16 | 2518 | 0.62 | 0.7 | -12345.33 | -11113.17 | [35] |
| Sediment | natural | 12 | 3796 | 0.77 | 0.47 | -16658.27 | -14665.48 | [35] |

**Figure S1.**
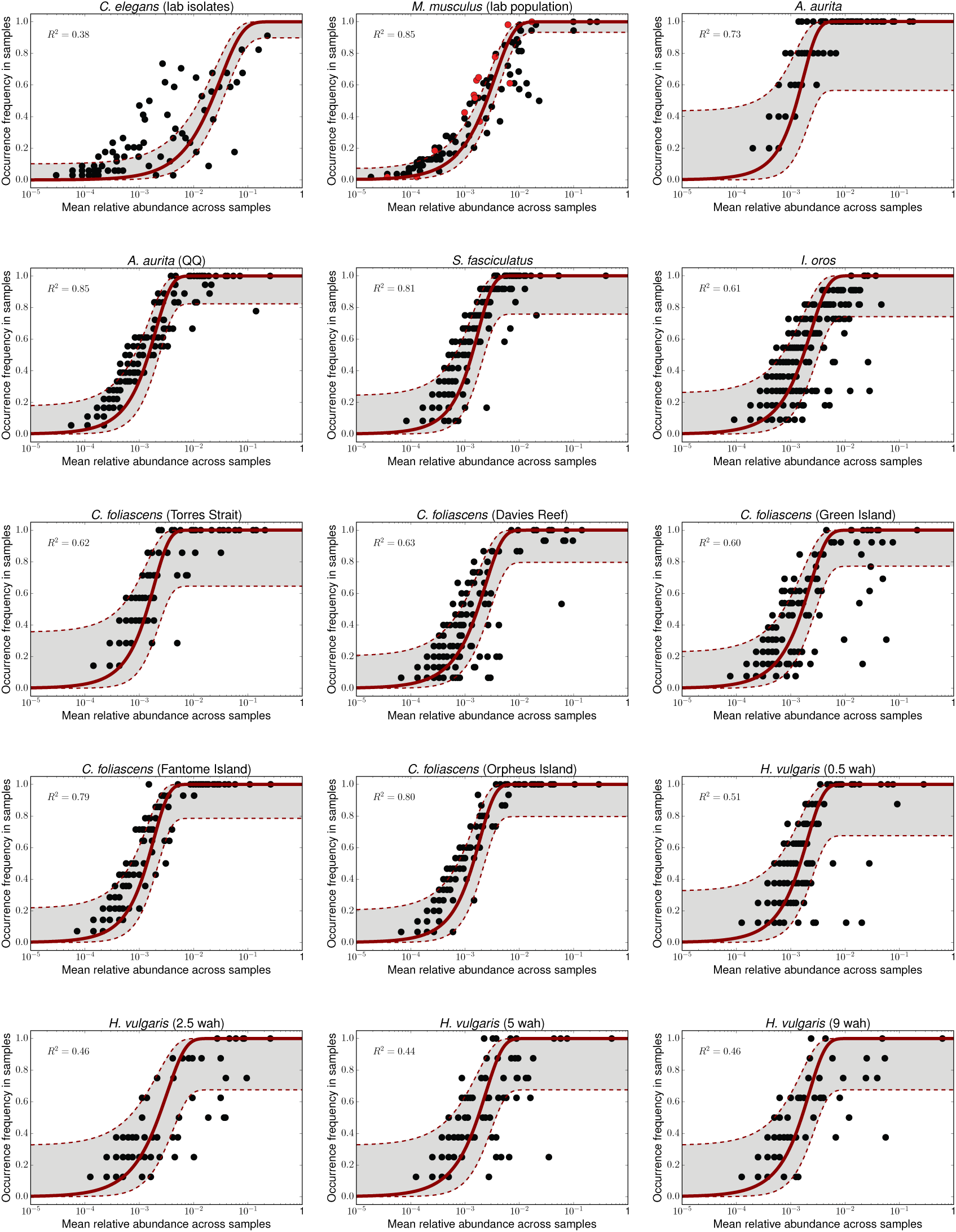
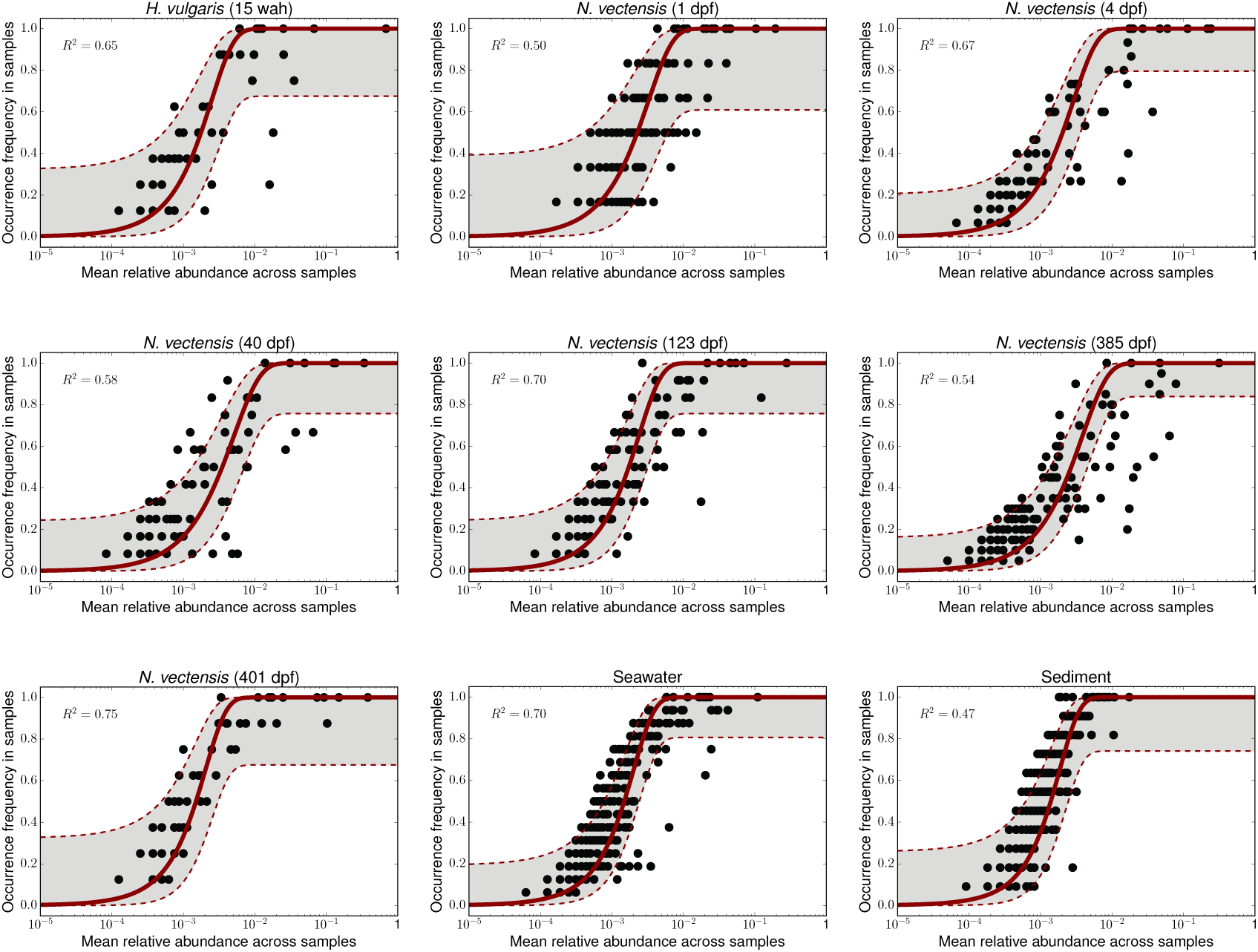
The relationship between mean relative abundance of taxa across all samples and the frequency with which they were detected in individual samples. Each dot represents a taxon and the solid line is the best-fitting neutral community expectation. The dashed lines and grey area depict the 95% confidence bands.

**Figure S2.**
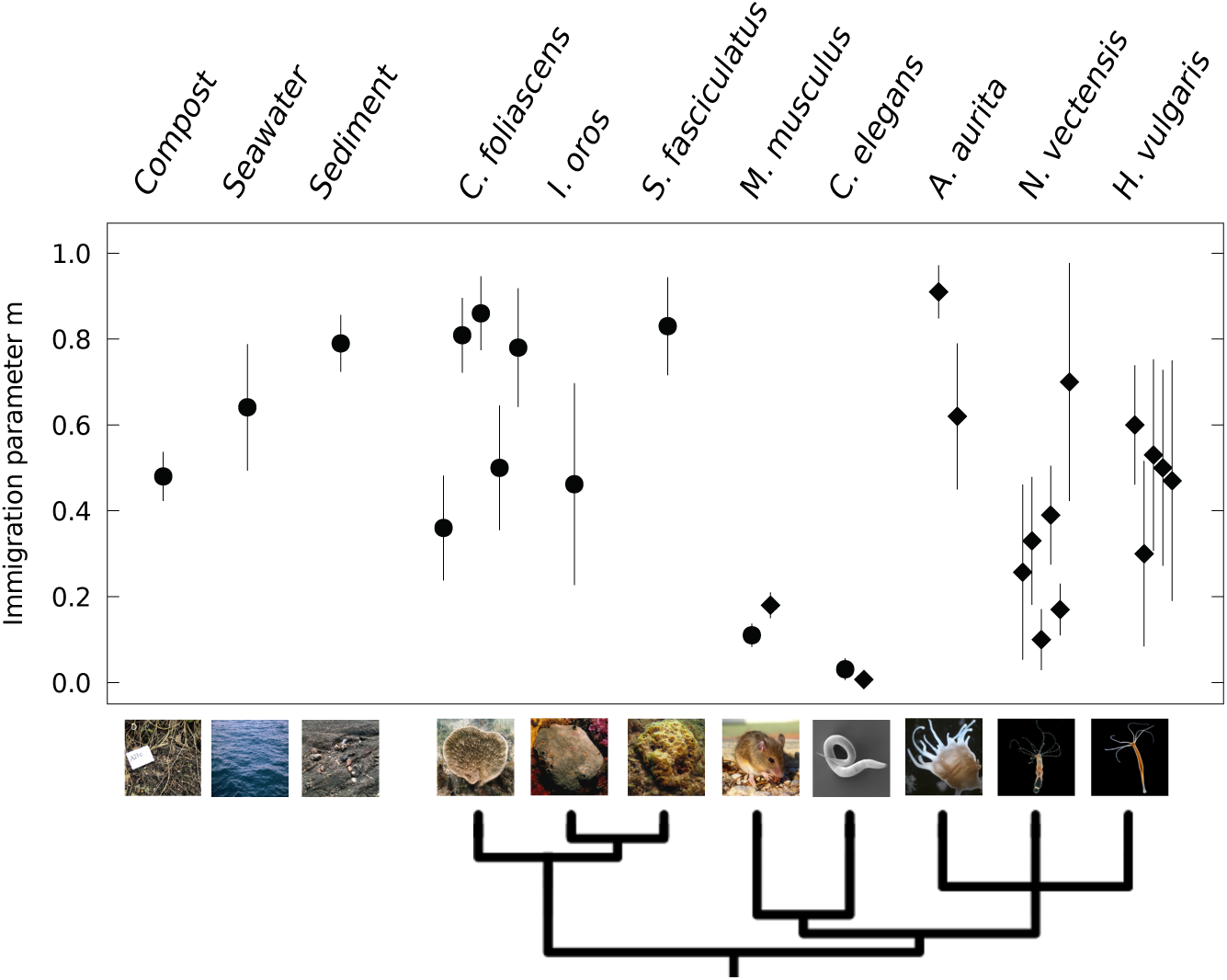
Overview of the best fit values for the immigration parameter *m* of the neutral model. Circles denote natural populations and diamonds denote laboratory populations, error bars indicate 95% bootstrap confidence intervals. The data for *C. foliascens* is from several different natural populations, while the data for *H. vulgaris* and *N. vectensis* is from different time points. Spread of points along the x-axis is added to increase visibility. Phylogeny generated with *phyloT* based on NCBI taxonomy. Sponge photographs courtesy of Susanna López-Legentil (UNC Wilmington) and Mari-Carmen Pineda (AIMS).

**Figure S3.**
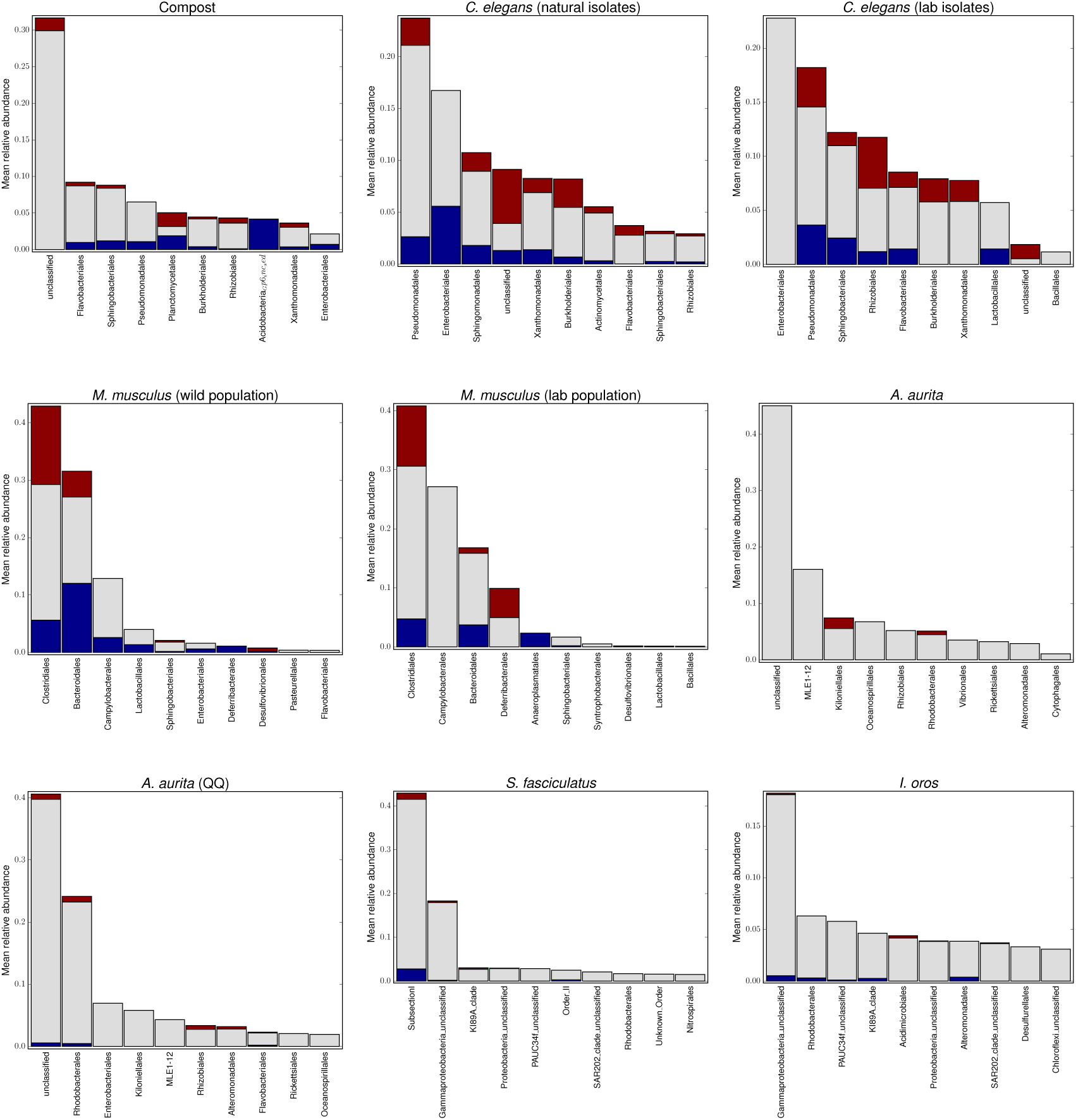
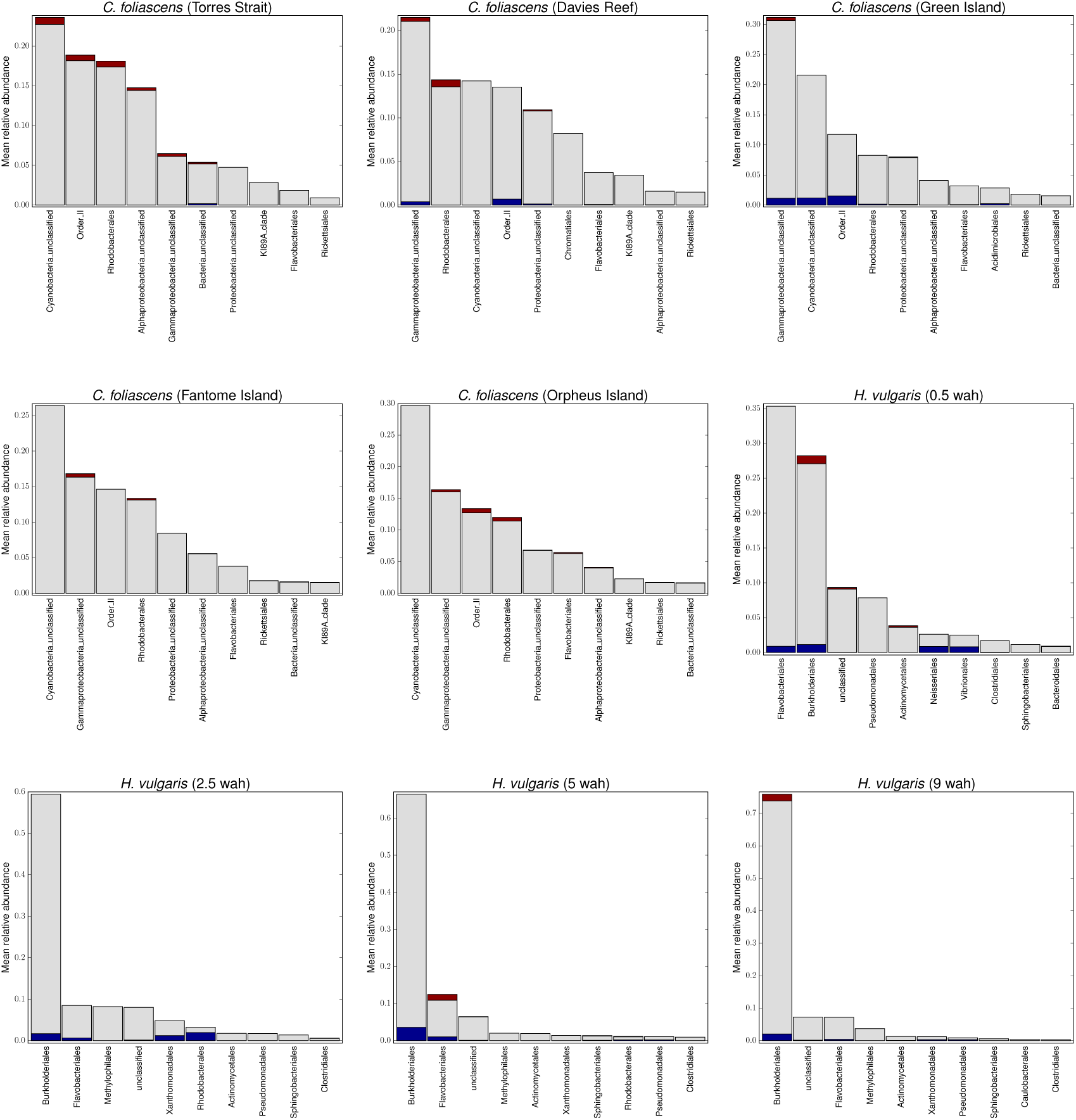
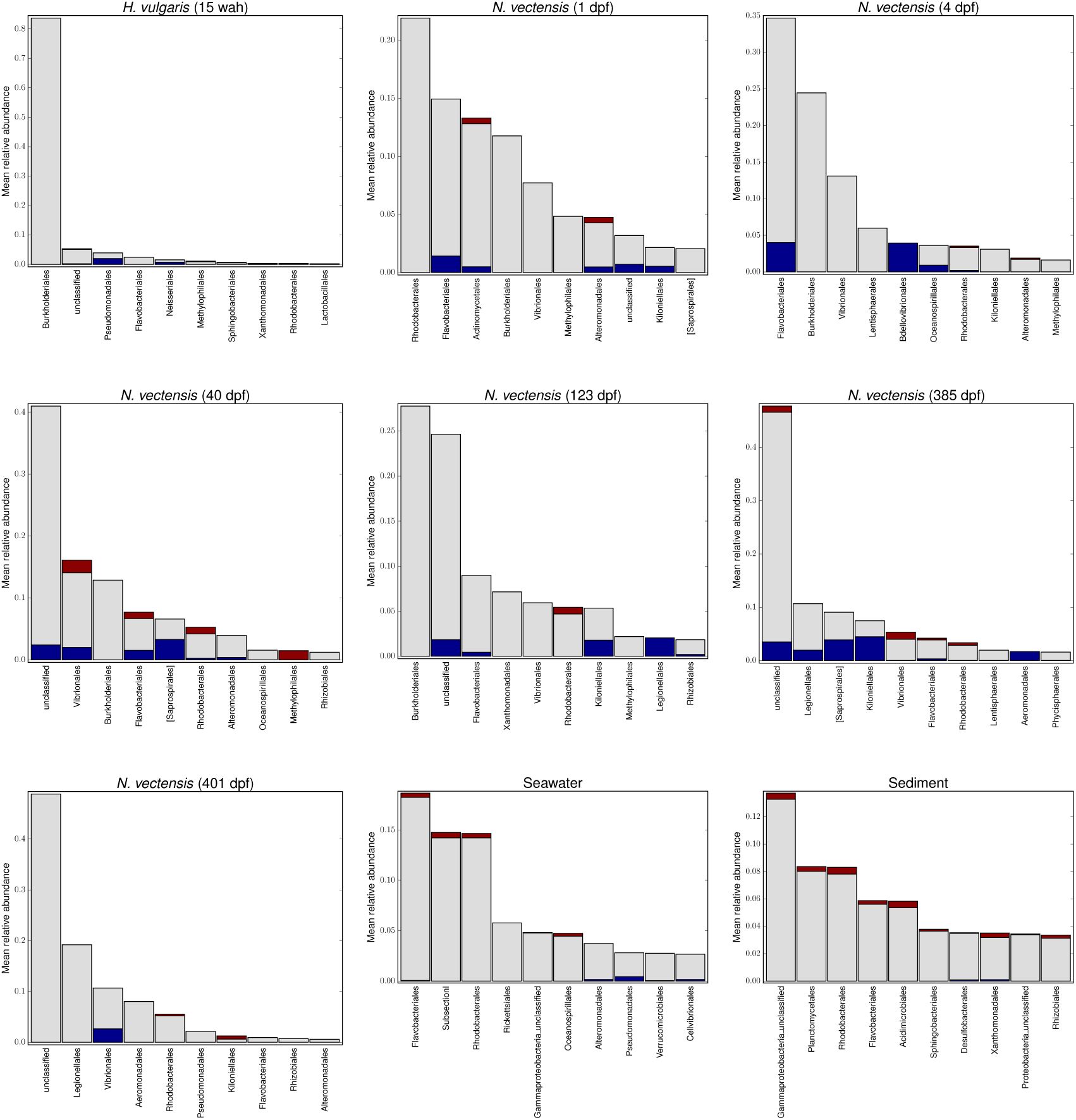
Bars show the mean relative abundance of the ten most abundant microbial orders in a specific host population. The colored sections indicate the fractions of OTUs within that order which were found above the neutral expectation (red), within the neutral expectation (grey) and below the neutral expectiation (blue).

**Table S2.** Non-neutral genera found in the *C. elegans* microbiota

| Domain | Phylum | Class | Order | Family | Genus |  |
| --- | --- | --- | --- | --- | --- | --- |
| Bacteria | Bacteroidetes | Flavobacteria | Flavobacteriales | Flavobacteriaceae | unclassified | over-represented |
| Bacteria | Proteobacteria | Alphaproteobacteria | Rhizobiales | Hyphomicrobiaceae | Devosia |  |
| Bacteria | Proteobacteria | Alphaproteobacteria | unclassified | unclassified | unclassified |  |
| Bacteria | Proteobacteria | Betaproteobacteria | Burkholderiales | Alcaligenaceae | unclassified |  |
| Bacteria | Proteobacteria | Betaproteobacteria | Burkholderiales | Oxalobacteraceae | unclassified |  |
| Bacteria | Proteobacteria | Betaproteobacteria | Burkholderiales | unclassified | unclassified |  |
| Bacteria | Proteobacteria | Betaproteobacteria | unclassified | unclassified | unclassified |  |
| Bacteria | Proteobacteria | Gammaproteobacteria | Pseudomonadales | Pseudomonadaceae | unclassified |  |
| Bacteria | Proteobacteria | Gammaproteobacteria | unclassified | unclassified | unclassified |  |
| Bacteria | Proteobacteria | Gammaproteobacteria | Xanthomonadales | Xanthomonadaceae | unclassified |  |
| Bacteria | unclassified | unclassified | unclassified | unclassified | unclassified |  |
| Bacteria | Proteobacteria | Alphaproteobacteria | Rhizobiales | Brucellaceae | Ochrobactrum | under-represented |

**Table S3.**
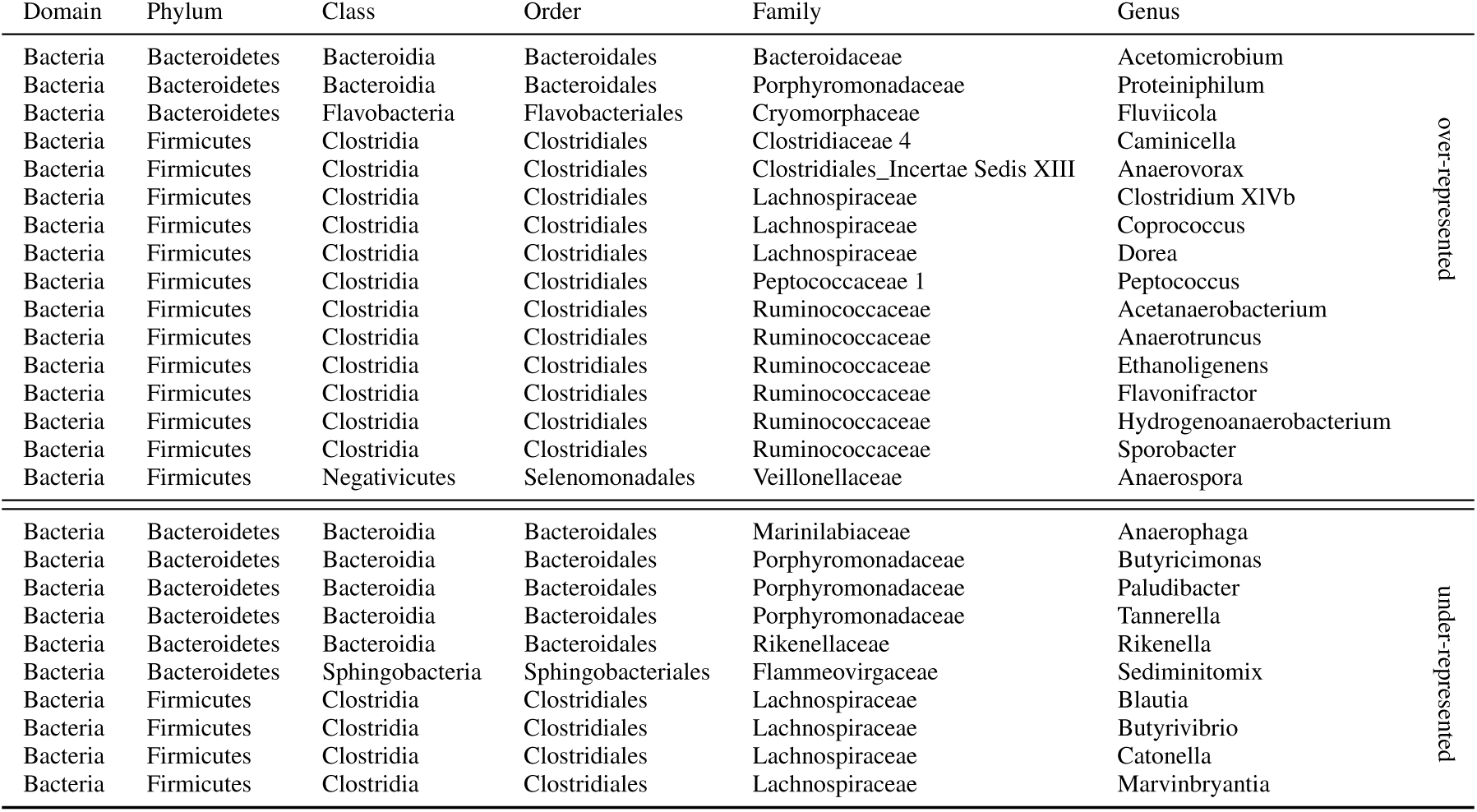
Non-neutral genera found in the *M. musculus* microbiota

**Figure S4.**
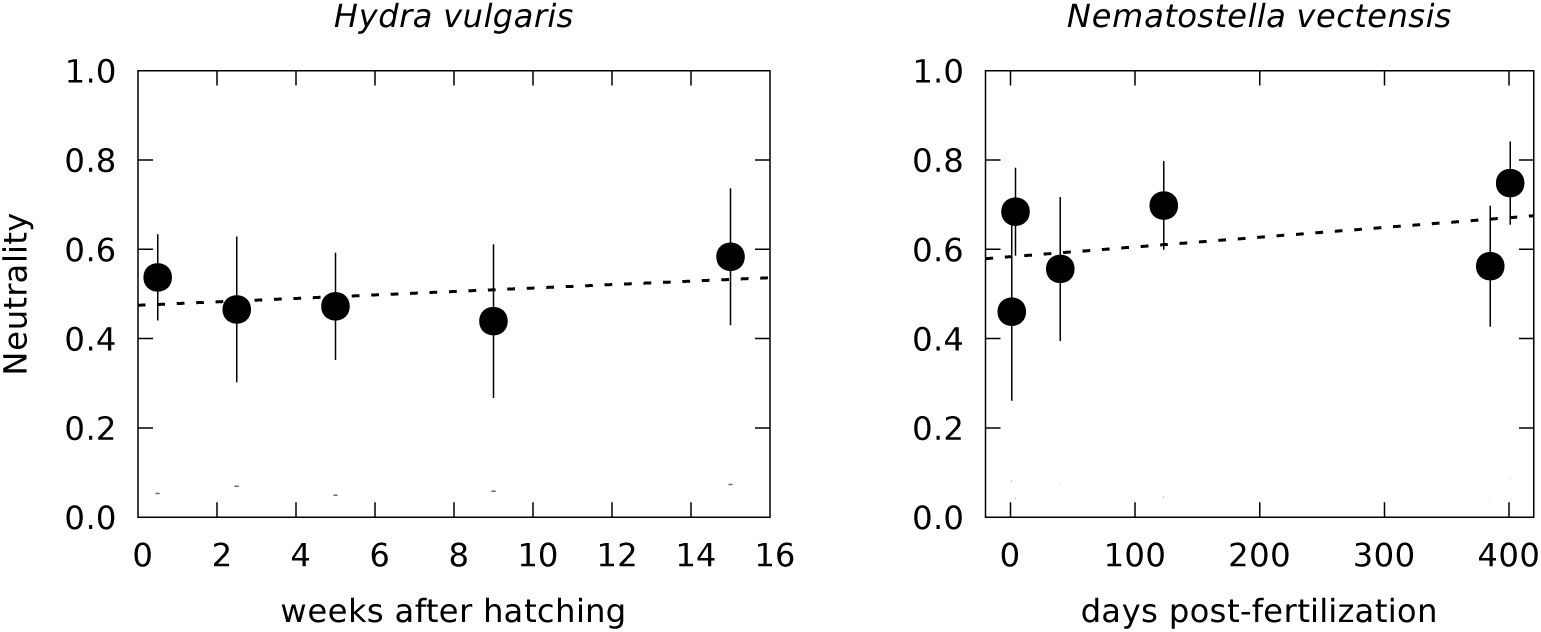
Neutrality of the microbiota of *Hydra vulgaris* and *Nematostella vectensis* at different time points during development and adult life stages (95% bootstrap confidence intervals attached). The dashed lines are the best linear fits, slopes are not significantly different from zero.

**Figure S5.**
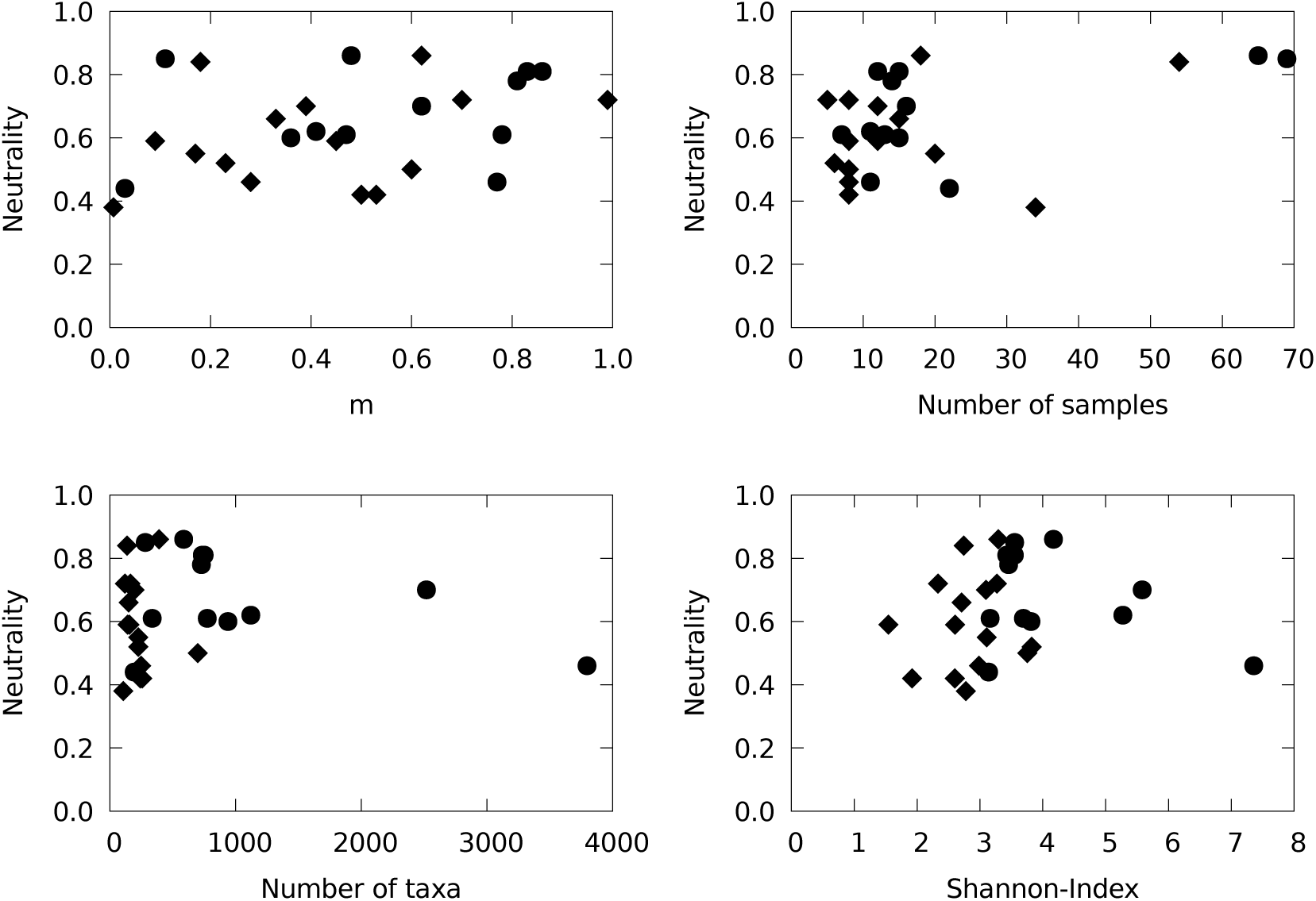
Consistency with the neutral model vs. the estimated dispersal parameter *m* (top left), the number of samples (top right), number of identified taxa (bottom left) and the Shannon-Index of diversity (bottom right). Circles denote natural populations and diamonds laboratory populations.

**Figure S6.**
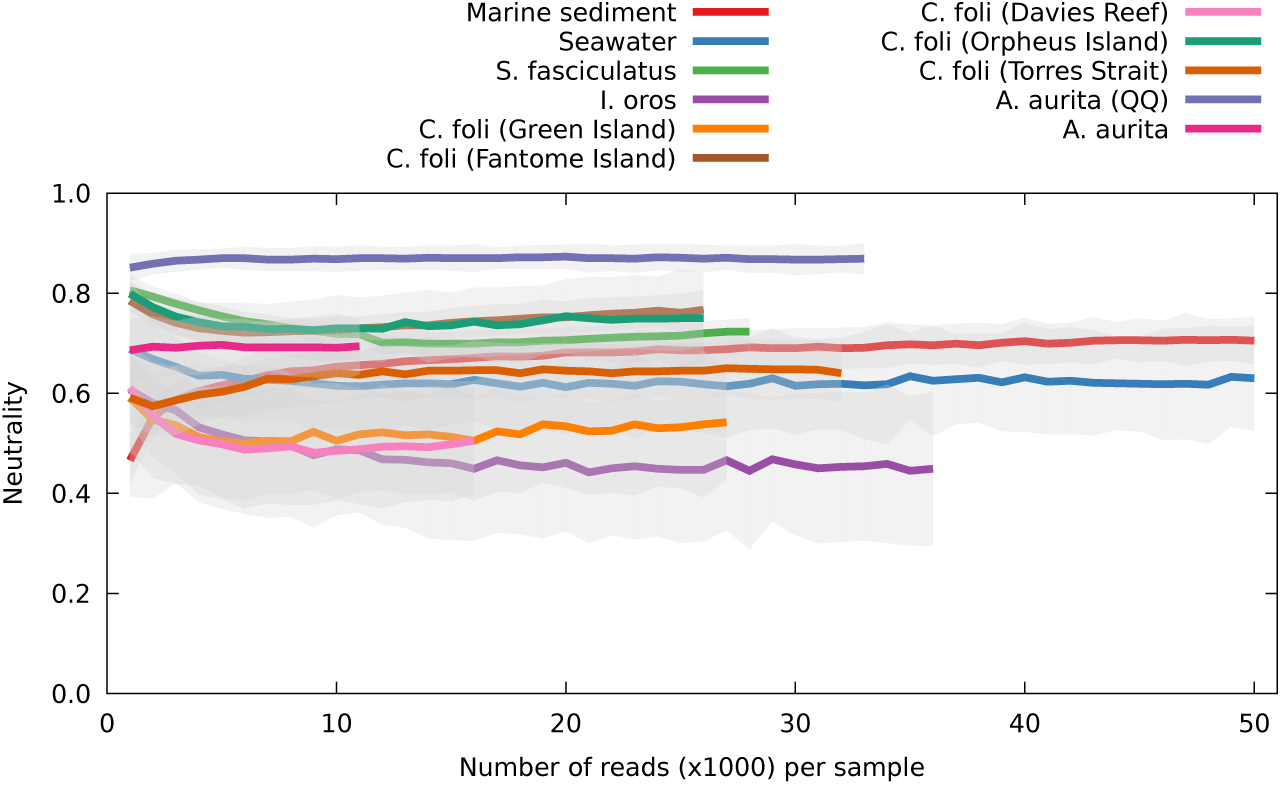
Impact of rarefaction on neutrality. Generally, consistency with the neutral model initially decreased with increasing read depth, until it leveled off as read depth increased further. For communities that showed a high consistency with the neutral model, varying read depths did not affect the results much, ranging from almost no effect at all for *A. aurita* to a slight drop in neutrality for the seawater samples from *R*^2^ = 0.7 at 1000 reads/sample to *R*^2^ ≈ 0.6 at 50000 reads/sample. Only for the communities associated with the sponge *I. oros* and two of the *C. foliascens* populations did read depth show a more pronounced effect, where in both cases neutrality dropped from *R*^2^ ≈ 0.6 at 1000 reads/sample to *R*^2^ ≈ 0.45 when read depth was exceeding 10000 reads/sample. Interestingly, an opposite trend was observed for the sediment samples, where neutrality increased from *R*^2^ ≈ 0.5 at 1000 reads/sample to *R*^2^ ≈ 0.7 at 10000 reads/sample. The minimal to moderate changes in neutrality with increasing read depth for some datasets potentially reflect the influence of rare, non-neutral taxa, which are only detected with higher read depths. Shaded areas indicate 95% bootstrap confidence intervals.

**Figure S7.**
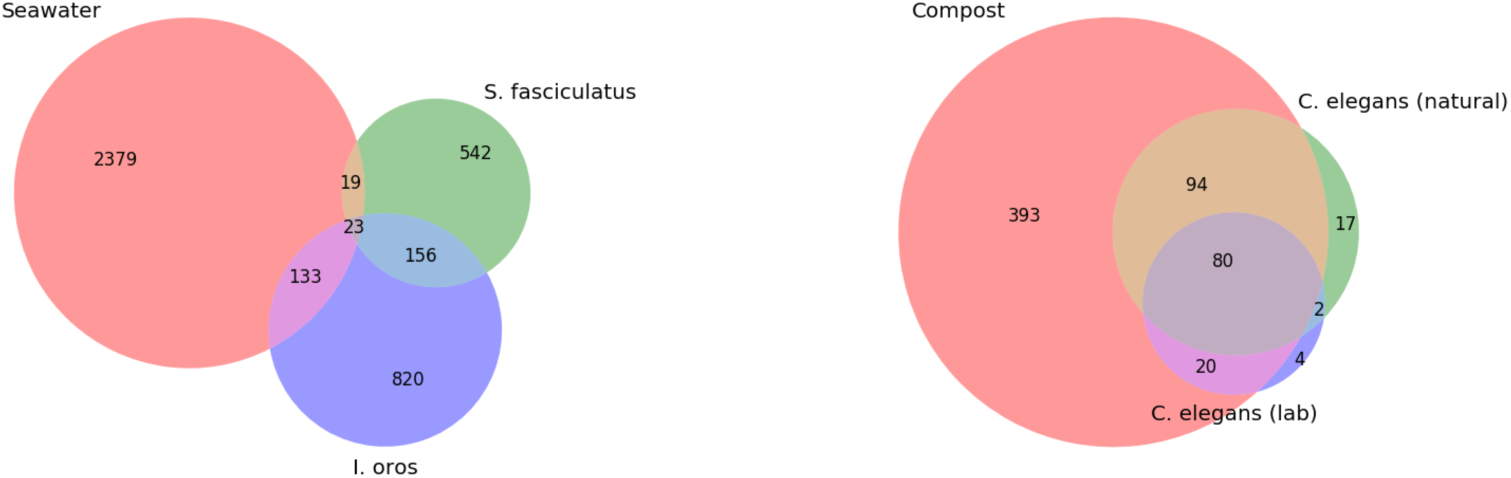
Overlap of taxa found in hosts and the environment. Left: For two sponge species, there is only a very small overlap between the sponge microbiota and the taxa found in seawater. Right: For *C. elegans*, only a subset of the environmentally available microbes is found in the worms.

## References

[1] Bäckhed F, Ding H, Wang T, Hooper LV, Koh GY, Nagy A, et al. The gut microbiota as an environmental factor that regulates fat storage. Proceedings of the National Academy of Sciences. 2004;101(44):15718–15723.

[2] Greiner T, Bäckhed F. Effects of the gut microbiota on obesity and glucose homeostasis. Trends in Endocrinology & Metabolism. 2011;22(4):117–123.

[3] Ridaura VK, Faith JJ, Rey FE, Cheng J, Duncan AE, Kau AL, et al. Gut microbiota from twins discordant for obesity modulate metabolism in mice. Science. 2013;341(6150):1241214.

[4] Belkaid Y, Hand TW. Role of the microbiota in immunity and inflammation. Cell. 2014;157(1):121–141.

[5] Bosch TC. Rethinking the role of immunity: lessons from *Hydra*. Trends in Immunology. 2014;35(10):495–502.

[6] Thaiss CA, Zmora N, Levy M, Elinav E. The microbiome and innate immunity. Nature. 2016;535(7610):65–74.

[7] Vuong HE, Yano JM, Fung TC, Hsiao EY. The microbiome and host behavior. Annual Review of Neuroscience. 2017;40:21–49.

[8] Murillo-Rincon AP, Klimovich A, Pemöller E, Taubenheim J, Mortzfeld B, Augustin R, et al. Spontaneous body contractions are modulated by the microbiome of *Hydra*. Scientific Reports. 2017;7(1):15937.

[9] Shapira M. Gut microbiotas and host evolution: scaling up symbiosis. Trends in Ecology & Evolution. 2016;31(7):539–549.

[10] Bosch TC, McFall-Ngai MJ. Metaorganisms as the new frontier. Zoology. 2011;114(4):185–190.

[11] McFall-Ngai M, Hadfield MG, Bosch TCG, Carey HV, Domazet-Lošo T, Douglas AE, et al. Animals in a bacterial world, a new imperative for the life sciences. Proceedings of the National Academy of Sciences. 2013;110(9):3229–3236.

[12] Sze MA, Schloss PD. Looking for a signal in the noise: revisiting obesity and the microbiome. mBio. 2016;7(4):e01018–16.

[13] Rothschild D, Weissbrod O, Barkan E, Kurilshikov A, Korem T, Zeevi D, et al. Environment dominates over host genetics in shaping human gut microbiota. Nature. 2018;555:210–215.

[14] Moran NA, Sloan DB. The hologenome concept: helpful or hollow? PLoS Biology. 2015;13(12):e1002311.

[15] Caswell H. Community Structure: A Neutral Model Analysis. Ecological Monographs. 1976;46(3):327–354.

[16] Hubbell SP. The unified neutral theory of biodiversity and biogeography. Monographs in Population Biology. Princeton, NJ, USA: Princeton University Press; 2001.

[17] Volkov I, Banavar JR, Hubbell SP, Maritan A. Neutral theory and relative species abundance in ecology. Nature. 2003;424(6952):1035–1037.

[18] Macarthur R, Wilson EO. The theory of island biogeography. Princeton University Press, Princeton, NJ; 1967.

[19] Rosindell J, Hubbell SP, He F, Harmon LJ, Etienne RS. The case for ecological neutral theory. Trends in Ecology & Evolution. 2012;27(4):203–208.

[20] Sloan WT, Lunn M, Woodcock S, Head IM, Nee S, Curtis TP. Quantifying the roles of immigration and chance in shaping prokaryote community structure. Environmental Microbiology. 2006;8(4):732–740.

[21] Woodcock S, Van Der Gast CJ, Bell T, Lunn M, Curtis TP, Head IM, et al. Neutral assembly of bacterial communities. FEMS Microbiology Ecology. 2007;62(2):171–180.

[22] Dumbrell AJ, Nelson M, Helgason T, Dytham C, Fitter AH. Relative roles of niche and neutral processes in structuring a soil microbial community. The ISME Journal. 2010;4(3):337–345.

[23] Ofiteru ID, Lunn M, Curtis TP, Wells GF, Criddle CS, Francis CA, et al. Combined niche and neutral effects in a microbial wastewater treatment community. Proceedings of the National Academy of Sciences. 2010;107(35):15345–15350.

[24] Lee JE, Buckley HL, Etienne RS, Lear G. Both species sorting and neutral processes drive assembly of bacterial communities in aquatic microcosms. FEMS Microbiology Ecology. 2013;86(2):288–302.

[25] Costello EK, Stagaman K, Dethlefsen L, Bohannan BJ, Relman DA. The application of ecological theory toward an understanding of the human microbiome. Science. 2012;336(6086):1255–1262.

[26] Sala C, Vitali S, Giampieri E, do Valle ÌF, Remondini D, Garagnani P, et al. Stochastic neutral modelling of the Gut Microbiota’s relative species abundance from next generation sequencing data. BMC Bioinformatics. 2016;17(Suppl 2):S16.

[27] Burns AR, Stephens WZ, Stagaman K, Wong S, Rawls JF, Guillemin K, et al. Contribution of neutral processes to the assembly of gut microbial communities in the zebrafish over host development. The ISME Journal. 2016;10(3):655–664.

[28] Li L, Ma ZS. Testing the neutral theory of biodiversity with human microbiome datasets. Scientific Reports. 2016;6:31448.

[29] Fisher CK, Mehta P. The transition between the niche and neutral regimes in ecology. Proceedings of the National Academy of Sciences. 2014;111(36):13111–13116.

[30] Kim HJ, Kim H, Kim JJ, Myeong NR, Kim T, Park T, et al. Fragile skin microbiomes in megacities are assembled by a predominantly niche-based process. Science Advances. 2018;4(3):e1701581.

[31] Venkataraman A, Bassis CM, Beck JM, Young VB, Curtis JL, Huffnagle GB, et al. Application of a neutral community model to assess structuring of the human lung microbiome. mBio. 2015;6(1):e02284–14.

[32] Adair KL, Wilson M, Bost A, Douglas AE. Microbial community assembly in wild populations of the fruit fly *Drosophila melanogaster*. The ISME Journal. 2018;.

[33] Adair KL, Douglas AE. Making a microbiome: the many determinants of host-associated microbial community composition. Current Opinion in Microbiology. 2017;35:23–29.

[34] Feuda R, Dohrmann M, Pett W, Philippe H, Rota-Stabelli O, Lartillot N, et al. Improved Modeling of Compositional Heterogeneity Supports Sponges as Sister to All Other Animals. Current Biology. 2017;27(24):3864–3870.

[35] Thomas T, Moitinho-Silva L, Lurgi M, Björk JR, Easson C, Astudillo-García C, et al. Diversity, structure and convergent evolution of the global sponge microbiome. Nature Communications. 2016;7:11870.

[36] Pita L, Rix L, Slaby BM, Franke A, Hentschel U. The sponge holobiont in a changing ocean: from microbes to ecosystems. Microbiome. 2018;6(1):46.

[37] Bosch, Thomas CG. Cnidarian-microbe interactions and the origin of innate immunity in metazoans. Annual Review of Microbiology. 2013;67:499–518.

[38] Samuel BS, Rowedder H, Braendle C, Félix MA, Ruvkun G. *Caenorhabditis elegans* responses to bacteria from its natural habitats. Proceedings of the National Academy of Sciences. 2016;113(27):E3941–E3949.

[39] Dirksen P, Marsh SA, Braker I, Heitland N, Wagner S, Nakad R, et al. The native microbiome of the nematode *Caenorhabditis elegans*: gateway to a new host-microbiome model. BMC Biology. 2016;14(1):38.

[40] Berg M, Stenuit B, Ho J, Wang A, Parke C, Knight M, et al. Assembly of the *Caenorhabditis elegans* gut microbiota from diverse soil microbial environments. The ISME Journal. 2016;10(8):1998–2009.

[41] Zhang F, Berg M, Dierking K, Félix MA, Shapira M, Samuel BS, et al. *Caenorhabditis elegans* as a model for microbiome research. Frontiers in Microbiology. 2017;8:485.

[42] Abolins S, King EC, Lazarou L, Weldon L, Hughes L, Drescher P, et al. The comparative immunology of wild and laboratory mice, *Mus musculus domesticus*. Nature Communications. 2017;8:14811.

[43] Linnenbrink M, Wang J, Hardouin EA, Künzel S, Metzler D, Baines JF. The role of biogeography in shaping diversity of the intestinal microbiota in house mice. Molecular Ecology. 2013;22(7):1904–1916.

[44] Wang J, Kalyan S, Steck N, Turner LM, Harr B, Künzel S, et al. Analysis of intestinal microbiota in hybrid house mice reveals evolutionary divergence in a vertebrate hologenome. Nature Communications. 2015;6:6440.

[45] Caporaso JG, Lauber CL, Walters WA, Berg-Lyons D, Huntley J, Fierer N, et al. Ultra-high-throughput microbial community analysis on the Illumina HiSeq and MiSeq platforms. The ISME Journal. 2012;6(8):1621–1624.

[46] Irazoqui JE, Urbach JM, Ausubel FM. Evolution of host innate defence: insights from *Caenorhabditis elegans* and primitive invertebrates. Nature Reviews Immunology. 2010;10(1):47–58.

[47] Müller WE, Müller IM. Origin of the metazoan immune system: Identification of the molecules and their functions in sponges. Integrative and Comparative Biology. 2003;43(2):281–292.

[48] Vega NM, Gore J. Stochastic assembly produces heterogeneous communities in the *Caenorhabditis elegans* intestine. PLoS Biology. 2017;15(3):e2000633.

[49] Flint HJ, Bayer EA, Rincon MT, Lamed R, White BA. Polysaccharide utilization by gut bacteria: potential for new insights from genomic analysis. Nature Reviews Microbiology. 2008;6(2):121–131.

[50] Daniel H, Gholami AM, Berry D, Desmarchelier C, Hahne H, Loh G, et al. High-fat diet alters gut microbiota physiology in mice. The ISME Journal. 2014;8(2):295–308.

[51] Fernandez MI, Regnault B, Mulet C, Tanguy M, Jay P, Sansonetti PJ, et al. Maturation of paneth cells induces the refractory state of newborn mice to *Shigella* infection. The Journal of Immunology. 2008;180(7):4924–4930.

[52] Hernandez-Agreda A, Gates RD, Ainsworth TD. Defining the core microbiome in corals’ microbial soup. Trends in Microbiology. 2017;25(2):125–140.

[53] Ermolaeva MA, Schumacher B. Insights from the worm: the *C. elegans* model for innate immunity. Seminars in Immunology. 2014;26(4):303–309.

[54] Dini-Andreote F, Stegen JC, van Elsas JD, Salles JF. Disentangling mechanisms that mediate the balance between stochastic and deterministic processes in microbial succession. Proceedings of the National Academy of Sciences. 2015;112(11):E1326–E1332.

[55] Mortzfeld BM, Urbanski S, Reitzel AM, Künzel S, Technau U, Fraune S. Response of bacterial colonization in *Nematostella vectensis* to development, environment and biogeography. Environmental Microbiology. 2016;18(6):1764–1781.

[56] Walker SC. When and why do nonneutral metacommunities appear neutral? Theoretical Population Biology. 2007;71(3):318–331.

[57] Mazzolini A, Gherardi M, Caselle M, Lagomarsino MC, Osella M. Statistics of shared components in complex component systems. Physical Review X. 2018;8(2):021023.

[58] De Wit R, Bouvier T. ‘*Everything is everywhere*, but, *the environment selects*’ what did Baas Becking and Beijerinck really say? Environmental Microbiology. 2006;8:755–758.

[59] Avershina E, Storrø O, Øien T, Johnsen R, Pope P, Rudi K. Major faecal microbiota shifts in composition and diversity with age in a geographically restricted cohort of mothers and their children. FEMS Microbiology Ecology. 2014;87(1):280–290.

[60] Franzenburg S, Walter J, Künzel S, Wang J, Baines JF, Bosch TCG, et al. Distinct antimicrobial peptide expression determines host species-specific bacterial associations. Proceedings of the National Academy of Sciences. 2013;110(39):E3730–E3738.

[61] Weiland-Bräuer N, Fischer MA, Pinnow N, Schmitz RA. Potential role of host-derived quorum quenching in modulating bacterial colonization in the moon jellyfish *Aurelia aurita*. Scientific Reports. 2019;9(1):34.

[62] Louca S, Parfrey LW, Doebeli M. Decoupling function and taxonomy in the global ocean microbiome. Science. 2016;353(6305):1272–1277.

[63] Louca S, Jacques SM, Pires AP, Leal JS, Srivastava DS, Parfrey LW, et al. High taxonomic variability despite stable functional structure across microbial communities. Nature Ecology & Evolution. 2016;1(1):0015.

[64] Lozupone CA, Stombaugh JI, Gordon JI, Jansson JK, Knight R. Diversity, stability and resilience of the human gut microbiota. Nature. 2012;489(7415):220–230.

[65] Louca S, Polz MF, Mazel F, Albright MB, Huber JA, O’Connor MI, et al. Function and functional redundancy in microbial systems. Nature ecology & evolution. 2018;p. 1.

[66] Coyte KZ, Schluter J, Foster KR. The ecology of the microbiome: Networks, competition, and stability. Science. 2015;350(6261):663–666.

[67] Fraune S, Anton-Erxleben F, Augustin R, Franzenburg S, Knop M, Schröder K, et al. Bacteria–bacteria interactions within the microbiota of the ancestral metazoan Hydra contribute to fungal resistance. The ISME Journal. 2015;9(7):1543.

[68] Kimura M. The Neutral Theory of Molecular Evolution. Cambridge, U.K.: Cambridge University Press; 1983.

[69] Franzenburg S, Fraune S, Altrock PM, Kuenzel S, Baines JF, Traulsen A, et al. Bacterial colonization of Hydra hatchlings follows a robust temporal pattern. ISME Journal. 2013;7:781–790.

[70] Newville M, Stensitzki T, Allen D, Ingargiola A. LMFIT: Non-Linear Least-Squares Minimization and Curve-Fitting for Python; 2014. Zenodo.

